# Divergent Pathways of Surfactant Protein C Maturation for Disease-Associated Isoforms

**DOI:** 10.1101/2025.10.03.679829

**Authors:** Sarah Bui, Anamarie Reineberg, Dakota Jones, Cheng-Lun Na, Joseph Kitzmiller, Luis R. Rodriguez, Aditi Murthy, Swati Iyer, Charlotte Cooper, Rea Chroneos, Yaniv Tomer, Surafel Mulugeta, Timothy E. Weaver, Darrell N. Kotton, Konstantinos-Dionysios Alysandratos, Jeffrey A. Whitsett, Michael F. Beers

## Abstract

Surfactant Protein C (SP-C), a hydrophobic protein exclusively synthesized and secreted by alveolar type II (AT2) cells, is important for reducing alveolar surface tension in the distal lung. Chronic interstitial pulmonary diseases have been associated with *SFTPC* mutations. However, a detailed understanding of SP-C maturation in the secretory pathway and disruptions caused by mutations has remained incomplete. The goal of this study was to comprehensively ascertain differences in trafficking and post-translational processing between wild-type and disease-associated SP-C mutants using doxycycline-inducible mouse lung epithelial (MLE-12) cell lines expressing either wildtype SP-C or the common clinical variant SP-C^I73T^, validated using primary AT2 cells isolated from a murine SP-C^I73T^ pulmonary fibrosis model and induced pluripotent stem cell (iPSC)-derived human alveolar type 2 cells (iAT2s) expressing the same mutant. In all 3 models SP-C^WT^ was highly concentrated in acidic LROs while SP-C^I73T^ accumulated on the plasma membrane, which was corroborated by inhibition of clathrin-mediated endocytosis, surface biotinylation, immunogold EM, immunofluorescent staining in non-permeabilized cells, and proteinase K protection assays supporting divergence of SP-C^I73T^ trafficking from SP-C^WT^. The exclusion of SP-C^I73T^ from normal routing occurred early in the biosynthetic pathway as Brefeldin A blocked processing of both SP-C proproteins, while a 20°C temperature shift caused selective accumulation of a processed proSP-C^WT^ intermediate, suggesting initial C-terminal cleavage of proSP-C^WT^ occurs in late-Golgi/ trans-Golgi network (TGN). This cleavage event was sensitive to DC1, an inhibitor of furin-related subtilisin-like proprotein convertase (PPC) family members. Site-directed mutagenesis of canonical residues K160, R167 within a predicted PPC recognition site in the proSP-C BRICHOS domain blocked its processing. Expression constructs encoding inhibitory pre-proprotein (pp) peptide fragments of Furin and ppPC7 each inhibited cleavage of proSP-C^WT^ by MLE-12 cells. Collectively, our data demonstrate that trafficking pathways for maturation of WT and mutant I73T SP-C diverge prior to the TGN where initial cleavage of the COOH-terminal SP-C propeptide occurs via a Furin-like proprotein convertase.

## INTRODUCTION

Surfactant Protein C (SP-C) is a small, hydrophobic peptide shown to be essential for reducing alveolar surface tension and maintaining normal lung compliance (1-3). SP-C is synthesized exclusively by alveolar type II epithelial (AT2) cells as a 21 kDa integral membrane precursor (proSP-C) that undergoes a series of tightly regulated post-translational modifications and proteolytic cleavages before being packaged into the lamellar body (LB), a lysosome related organelle (LRO), for release into the alveolar lining fluid by regulated exocytosis (1, 4-7). The native proSP-C is a type 2 bitopic transmembrane protein comprised of four domains: 1) an NH2 terminal cytosolic domain required for post-Golgi targeting; 2) the mature SP-C protein which serves as the membrane anchor transmembrane helix; 3) an unstructured linker domain in the proximal COOH propeptide; 4) a distal COOH-terminal BRICHOS domain that contributes to proprotein folding (8-12). Defects in SP-C biosynthesis from autosomal dominant mutations in the COOH propeptide regions encoded by the *SFTPC* gene can impair protein folding, trafficking, or proteolytic maturation and underlie a subset of familial interstitial lung diseases (ILD) in children and adults (13-19).

The majority of published data on SP-C biology has supported a model wherein maturation of wild-type proSP-C occurs via a classical, anterograde trafficking pathway progressing from Golgi to late endosomes, multivesicular bodies (MVB) and ultimately LB, which generates the 3.7 kDa secreted form (9, 20-23). In contrast, mutant isoforms of SP-C segregate into 2 major functional classes: (i) Aggregation-prone mutants within the BRICHOS domain do not traffic beyond the endoplasmic reticulum (ER) because they are unable to fold correctly and elicit endoplasmic reticulum (ER) stress and AT2 cell apoptosis (24-26); (ii) “Trafficking” mutants, such as the common pathogenic variant SP-C^I73T^, are misrouted to the plasma membrane, cause endolysosomal stress, inhibit macroautophagy, and alter mitophagy (27, 28). Proof-of-concept studies *in vivo* using knock-in murine models expressing either SP-C^I73T^ or a representative BRICHOS misfolding mutation (SP-C^C121G^; SP-C^C185G^) each demonstrate the development of a spontaneous lung fibrosis (29-32).

While the latter stages of proSP-C processing have been well characterized, particularly NH2-terminal remodeling events mediated by two proteolytic cleavages involving Cathepsin H (33) and an aspartic protease (Napsin A or Progastricsin C) occurring in late endosomal or LB compartment (34), more proximal post-translational processing events involving COOH propeptide remodeling as well as the impact of *SFTPC* mutations on trafficking remain incompletely defined. In particular, the initial cleavage event removing the distal COOH-terminal propeptide (which precedes the aforementioned N-terminal cleavage events), has not been attributed to a specific enzyme or localized to a defined subcellular compartment. Similarly, it remains controversial whether routing of wild-type SP-C and the SP-C^I73T^ variant overlap (22, 23, 35). Thus, a clearer understanding of trafficking and post-translational processing of SP-C and its mutants is essential to constructing a complete model of SP-C biogenesis that can also provide insights into key vulnerabilities in disease-associated *SFTPC* mutants that drive familial ILD pathogenesis.

To address these gaps in knowledge, we investigated SP-C biosynthesis to comprehensively ascertain similarities and differences in post-translational processing between wild-type and disease-associated *SFTPC* mutants, focusing on intracellular trafficking itineraries and early proteolytic events that occur prior to lamellar body localization. Our study used a multi-model approach consisting of a combination of doxycycline-inducible lung epithelial cell lines, primary murine AT2 cells, and human iPSC-derived AT2 cells (iAT2s) subjected to pharmacologic perturbations, biochemical fractionation, and site-directed mutagenesis. In doing so, we add to the evidence that SP-C follows a classical anterograde pathway from ER through to LB, but that pathogenic mutants diverge from this course early. We also identify the compartment and candidate enzymes responsible for the initial proteolytic processing event.

## MATERIALS AND METHODS

### Mouse model of pulmonary fibrosis

Tamoxifen-inducible Sftpc^I73T/I73T^ Rosa26ERT2FlpO^+/+^ (a.k.a. SP-C^I73T^) mice expressing an NH_2_-terminal HA-tagged murine Sftpc^I73T^ mutant allele in the endogenous mouse Sftpc locus were previously generated as reported (29). The SftpcI73T knock-in allele bearing 218 T>C point mutation (a.k.a. SP-C^I73T^ ^KI/KI^) was generated by Timothy Weaver using the same strategy previously described (36).

### Cell Culture

The parental murine lung epithelial cell line MLE-12 (CRL-2110) (37) was originally obtained from the American Type Culture Collection (Manassas, VA) and maintained in culture at 37 °C and 5% CO_2_ in HITES medium [Ham’s F-12 medium (50:50 mixture) containing 0.005 mg/ml insulin, 0.01 mg/ml transferrin, 30 nM sodium selenite, 10 nM hydrocortisone, 10 nM β-estradiol, 10 mM HEPES buffer, and 2 mM l-glutamine] supplemented with 2% fetal bovine serum and antibiotics. These cells contain organelles with some of the ultrastructural characteristics of lamellar bodies, and secrete phospholipids.

### Stable Doxycycline-inducible GFP-SP-C cell lines preparation by lentivirus

The subcloning of human SP-C (*SFTPC*^WT^), BRICHOS mutant SP-C (*SFTPC*^C121G^), and I73T mutant SP-C (*SFTPC*^I73T^) into the pEGFP-C1 and pcDNA3 expression vectors was previously described (38, 39). Generation of doxycycline inducible MLE-12s was performed using lentiviral infection followed by clonal selection. Generation of transfer plasmid was performed by subcloning GFP-SP-C locus into backbone plasmid pCW57.1 (pCW57.1 was a gift from David Root, Addgene plasmid # 41393). Lentiviral particles were packaged in HEK293T cells via lipofectamine 2000 transfection with envelope plasmid pMD2.G and packaging plasmid psPAX2 (both gifts from Didier Trono, Addgene plasmids #12259 and # 12260). MLE-12 cells were infected for 24 hours followed by puromycin selection to enrich for infected cells. Individual clones were expanded and screened for expression of SP-C proprotein.

### Reagents and Materials

Except where noted, all other reagents were electrophoretic or immunological grade and purchased from either Millipore Sigma (St. Louis, MO) or Thermo-Fisher (Pittsburgh, PA). Antibodies used for flow-cytometry, FACS, and immunoblotting were obtained either from commercial sources or generated in-house and validated as previously published and are summarized in **Supplemental Table 1**.

### Lectin Staining

MLE-12 cells transfected with EGFP-SP-C constructs were seeded on poly-l-lysine coated glass coverslips. The following day, double immunofluorescence staining with cell surface Wheat Germ Agglutin (WGA) and an epitope specific polyclonal antibody directed against the proSP-C COOH-terminal (40) was performed without permeabilization as previously described (22). In brief, intact cells were incubated with ice-cold PBS for 15 min and cells were incubated with CTerm proSP-C antibody for 1h on ice. Cells were washed in PBS and further incubated in TRITC-conjugated WGA (2 μg/ml) (Vector Laboratories) for 30 min on ice. Cells were washed with cold PBS, followed by fixation with 1% PFA for 15 min prior to secondary antibody staining and Hoechst for nuclear staining before mounting the coverslips with Mowiol. Fluorescence images were taken using a 60x oil objective on the Nikon Eclipse Ti2 and processed with extended depth of focus (EDF) projection. To quantify the localization of proSP-C relative to the plasma membrane in these images, relative fluorescence intensity was plotted along a random line through the region using Nikon NIS-Elements AR analysis software.

### Co-localization of EGFP-SP-C with Lysotracker and ER Tracker in Live Cells

MLE-12 cells were seeded in Nunc Glass Bottom Dishes (Φ 12 mm, Thermo Fisher Scientific, Inc.) at a density of 1.5–2.0 × 10^4^ per well in 2% Hites Media. After overnight doxycycline-induction of GFP-SP-C, the cells were washed with HBSS and incubated with 50 nM lysotracker Red DND-99 (Thermo Fisher Scientific) or 100 nM ER-Tracker™ Red (Thermo Fisher Scientific) diluted in pre-warmed OPTI-MEM for 30 min in a 5% CO2 at 37 °C. The supernatant was discarded, rinsed three times with probe-free HBSS, and the cells were post-incubated with pre-warmed phenol-free imaging medium for live-cell fluorescent imaging at 60× on the Nikon Eclipse Ti2 and processed with extended depth of focus (EDF) projection.

### Immunofluorescence of MLE-12 cells

Cells grown on 1.5 circular glass coverslips were washed with PBS and fixed in 4% paraformaldehyde for 10 min at room temperature. Cells were permeabilized with 0.2% Triton-X in PBS for 15 min. The samples were blocked in 3% BSA (in 1X PBS) for 60 min at room temperature, followed by incubating with primary and secondary antibodies. The cell nucleus was stained using Hoechst 33258 (Sigma-Aldrich). Coverslips were mounted in Permount™ Mounting Medium (ThermoFisher) and cured for 24 h before imaging. The following primary antibodies were used for immunostaining in this study: (**Supplemental Table 1**). Imaging was performed on a Nikon Eclipse Ti2 at 60× magnification. Image acquisition and deconvolution were performed with the Nikon software program. Images were further cropped or adjusted using ImageJ (National Institutes of Health).

### SDS/PAGE and Immunoblotting

SDS/PAGE and western immunoblotting of cell lysates was performed as described (41). In brief, harvested cell pellets were resuspended in lysis buffer (20 mM Tris-HCl pH 8.0, 150 mM NaCl, 1% Triton X-100 supplemented with protease inhibitor cocktail (Thermo Fisher) and the lysate cleared via centrifugation at 20,000 *g* and 4°C for 10 min in a benchtop microcentrifuge. Protein quantification was performed using a Bradford Kit (Bio-Rad). Cell lysates were boiled in SDS loading buffer (6×, 300 mM Tris-HCl, pH 6.8, 6% SDS, 0.06% Bromophenol Blue, 36% (v/v) glycerol, 12 mM DTT) and proteins separated by SDS-PAGE and then transferred to nitrocellulose membranes using a semi-dry or wet transfer machine (BioRad). For immunoblotting, membranes were blocked in 5% milk in PBST (0.1% Tween 20 in PBS) and incubated in primary antibody followed by species specific HRP-conjugated secondary antibody. Antibodies used in this study are shown in **Supplemental Table 1**. Bands detected by enhanced chemiluminescence using SuperSignal WestPicoPlus substrate (ThermoFisher) were acquired by direct scanning using an LiCor Odyssey Fc Imaging Station (LI-COR Biotechnology, Lincoln, NE) and quantified using the manufacturers’ software or ImageJ.

### Transferrin Endocytosis Assay

MLE12 cells seeded on poly-l-lysine coated coverslips were washed with PBS and incubated for 20 mins in serum-free OPTI-MEM with or without Pitstop2 (20μM) before being pulsed with AlexaFluor-488 conjugated mouse-Transferrin (50μg/ml) in the same media for 30 mins at 37°C. Cells were quickly washed in PBS and immediately fixed in 2% paraformaldehyde. Hoechst 33258 (Sigma-Aldrich) was used to stain nuclear DNA. Fluorescence images were taken using a 60x oil objective on Nikon Eclipse Ti2 and processed with EDF Focused Image projection for each independent experiment to assess internalized Transferrin. All images were captured and processed with the same setting. Quantifications were performed to calculate sum intensity of selected ROIs using the Nikon NIS-Elements AR analysis software.

### Cell Surface Biotinylation

EGFP-SP-C expression in MLE-12 cells seeded on 15-cm dishes was induced overnight with doxycycline (2.5 μM), followed by 2 hours of Pitstop2 (20μM) or DMSO (vehicle control) in serum-free OPTI-MEM with doxycycline. Cells were washed twice with ice-cold PBS, treated with 10 ml of 0.5 mg/ml NHS-SS-biotin (Thermo Fisher) in PBS for 20 min in the cold room, and quenched by 100 mM glycine in PBS for 10 min. After three washes with ice-cold PBS, cells were lysed in lysis buffer (20 mM Tris-HCl, pH 8.0, 150 mM NaCl, 1% Triton X-100, 0.1% SDS, 0.1% sodium deoxycholate, 1 mM EDTA, 50 mM sodium fluoride, 20 mM sodium orthovanadate, and 1× protease inhibitor cocktail). After centrifugation, the supernatants were adjusted to the same concentration and incubated with streptavidin-agarose beads overnight. After extensive washing, beads were boiled in SDS loading buffer with 40 mM DTT. Proteins were separated by SDS-PAGE and analyzed by immunoblotting as described.

### Proteinase K Protection Assay

Expression of wildtype and mutant GFP-SFTPC isoforms was induced in MLE12 lines with doxycycline overnight and treated with Pitstop2 (20μM) for 20 minutes the following day. Cells were then washed with PBS and incubated in pre-warmed PBS with calcium/magnesium containing 1 μg/mL Proteinase K (Sigma p7850) for 10 minutes at 37°C. Positive control (with Triton-X permeabilization before Proteinase K incubation) and negative control conditions (with neither Triton-X nor Proteinase K), were performed in parallel. Proteinase K was then inhibited by adding 5 mM PMSF to the treated cells, incubated on ice for 10 min, before proceeding to cell lysis.

### iPSC line generation and maintenance

All experiments involving the differentiation of human iPSC lines were performed with the approval of the Institutional Review Board of Boston University (protocol H-33122). The SPC2 iPSC line clones SPC2-ST-C11 and SPC2-ST-B2 used in this study were previously generated using TALENs to insert a tdTomato fluorescent reporter at the translation initiation (ATG) site of the endogenous SFTPC locus of the parental SPC2 iPSC line, resulting in the generation of either corrected (SPC2-ST-B2 clone; SFTPCtdT/WT) or mutant (SPC2-ST-C11 clone; SFTPCI73T/tdT) iPSC clones (23). iPSCs used in this study demonstrated a normal karyotype when analyzed by G-banding and/or array Comparative Genomic Hybridization (aCGH, Cell Line Genetics). iPSCs were maintained in feeder-free conditions on growth factor-reduced Matrigel (Corning) in 6-well tissue culture dishes (Corning) in mTeSR1 media (StemCell Technologies), using gentle cell dissociation reagent for passaging. Further details of iPSC derivation, characterization, and culture are available for free download at https://crem.bu.edu/cores-protocols/#protocols.

### iPSC-directed Differentiation into Alveolar Epithelial Type 2 Cells (iAT2s)

To generate iAT2s, PSC-directed differentiation via definitive endoderm into NKX2-1 lung progenitors was performed using methods we have previously described (23, 42, 43). On days 15-17 of differentiation, live cells were sorted on a high-speed cell sorter (MoFlo Astrios EQ) to isolate NKX2-1+ lung progenitors based on CD47hiCD26− gating (43). Sorted lung progenitors were resuspended in undiluted growth factor-reduced 3D Matrigel (Corning) at a density of 400 cells/μl, and distal/alveolar differentiation of cells was performed in CK+DCI medium, consisting of complete serum-free differentiation medium (cSFDM) base supplemented with 3 μM CHIR99021, 10 ng/mL recombinant human KGF (CK), 50 nM dexamethasone (Sigma), 0.1 mM 8-Bromoadenosine 3′,5′-cyclic monophosphate sodium salt (Sigma), and 0.1 mM 3-Isobutyl-1-methylxanthine (IBMX; Sigma) (DCI). The resulting epithelial spheres were passaged without further sorting on approximate day 30 (day 28-32) of differentiation, and a brief period (4-5 days) of CHIR99021 withdrawal followed by one week of CHIR99021 addback was performed to achieve iAT2 maturation, as previously shown (23). After this 2-week period, SFTPCtdTomato+ cells were purified by fluorescence activated cell sorting (FACS) to establish pure cultures of iAT2s. iAT2s were then maintained through serial passaging as self-renewing monolayered epithelial spheres (“alveolospheres”) by plating in 3D Matrigel (Corning) droplets at a density of 400 cells/μl with refeeding every other day with CK+DCI medium, according to our published protocol (44). iAT2 culture quality and purity were monitored at each passage by flow cytometry, with > 90% of cells expressing SFTPCtdTomato over time, as we have previously detailed (23, 44).

Established alveolospheres were treated with vehicle control, Pitstop 2 (10μM) for 2 hours, or Bafilomycin A1 (50nM) for 16 hours. For cryo-embedding, alveolospheres were fixed in 4% paraformaldehyde (PFA) for 20 minutes at room temperature. Samples were dehydrated in sucrose, frozen in OCT, and cryosectioned at 6 µm thickness. Slides were blocked for 1 hr at RT in a humid chamber with blocking buffer ([1% BSA [Sigma], 0.1% Triton X-100 [Fisher BioReagents] in PBS). The slides were then stained in blocking buffer overnight at 4°C with primary antibodies (**Supplemental Table 1**). The following day, the slides were washed three times for 5 min while gently shaking at RT with PBS and stained for 45 min in blocking buffer with secondary antibodies (**Supplemental Table 1**). Slides were then washed three times for 5 min while gently shaking at RT with PBS and stained with Hoechst (1:3,000 dilution, Thermo Fisher Scientific), for 5 min and washed in PBS as mentioned above. Slides were mounted with ProLong Gold antifade reagent, (P36930, Thermo Fisher Scientific) mounting medium with Number 1.5 coverslip. Images were acquired and processed with LAS X software on the Leica Stellaris 5 Confocal. Whole cell lysates from Pitstop 2 or Bafilomycin A1 treated alveolospheres were collected in parallel and analysed via Western blot as described in prior section (see **SDS/PAGE and Immunoblotting; Supplemental Table 1**).

### Electron Microscopy

Preparation of lung tissue and acquisition of TEM images of lung sections was performed in the Electron Microscopy Resource Laboratory in the Perelman School of Medicine based on the method of Hayat that includes postfixation in 2.0% osmium tetroxide with 1.5% potassium ferricyanide. Cut thin sections (60–80 nm) were stained in situ on copper grids with uranyl acetate and lead citrate and examined with a JEOL 1010 electron microscope fitted with a Hamamatsu digital camera and AMT Advantage image capture software.

### Immuno EM

Six to eight week old SP-C^wt/wt^ and SP-C^I73T^ ^Ki/KI^ mouse lungs were fixed by tracheal instillation with 4% paraformaldehyde-lysine-sodium periodate, 0.1% glutaraldehyde, and 0.1% CaCl2 in 0.2M HEPES, pH 7.2, cryo protected, and processed for immunogold as described previously (45, 46). Localization of Pro-SP-C to the AT2 cells of SP-C^wt/wt^ and SP-C^I73T^ ^KI/KI^ mice was demonstrated by incubating 100 nm ultrathin mouse lung frozen sections with rabbit antisera directed against the N-terminus of Pro-SP-C, followed by visualization with 10 nm protein A gold. Electron micrographs were acquired using a Hitachi H-7650 transmission electron microscope (Hitachi High Technologies America) equipped with an AMT CCD camera (Advanced Microscopy Techniques).

### Histology and Immunofluorescence of Lung Tissues

Lung tissue was from unidentified normal donors provided by the BRINDL program at the University of Rochester. Tissue was obtained from a deidentified donor bearing an SFTPC^I73T^ mutation at lung transplant provided by William Gower at the University of North Carolina, Chapel Hill, NC with permission. Human lung tissues were fixed in 10% formalin and embedded in paraffin. Sections were melted at 60°C for two hours and rehydrated through xylene and alcohol, and finally in PBS. Antigen retrieval was performed in 0.1 M citrate buffer (pH 6.0) by microwaving. Slides were blocked for 2 hours at room temperature using 4% normal donkey serum (Jackson Immuno Research Laboratories) in PBS containing 0.2% Triton X-100, and then incubated with primary antibodies diluted in blocking buffer for approximately 16 hours at 4°C. For immunofluorescence, slides were blocked for 2 hours at room temperature using 4% normal donkey serum (Jackson Immuno Research Laboratories) in PBS containing 0.2% Triton X-100, and then incubated with primary antibodies diluted in blocking buffer for approximately 16 hours at 4°C. Primary antibodies included ABCA3 (1:100, Seven Hills Bioreagents), and SFTPC (1:250, Seven Hills Bioreagents). Appropriate secondary antibodies conjugated to Alexa Fluor 488, Alexa Fluor 568, or Alexa Fluor 633/647 (Thermo Fisher Scientific, Jackson Immuno Research) were used at a dilution of 1:200 in blocking buffer for 1 hour at room temperature. Nuclei were counterstained with DAPI (1 μg/ml), (D21490, Thermo Fisher Scientific). Sections were mounted using ProLong Gold antifade reagent, (P36930, Thermo Fisher Scientific) mounting medium with Number 1.5 coverslip. Tissue sections stained by immunofluorescence were imaged on an inverted Nikon AXR confocal microscope at 100X super resolution magnification at 0.03µm/px a NA 1.27 objective and using a 1.2 AU pinhole. Image acquisition and deconvolution were performed with Nikon Elements software. 3D images were exported using Imaris (Bitplane) software.

### Isolation of Golgi Membrane Fractions by Sucrose Gradient Centrifugation

Golgi membrane fractions were isolated using published methods (47). Cells from four 15 cm² cell culture dishes were harvested with PBS containing 0.5x protease and phosphatase inhibitors (1.2 ml per flask). After centrifugation for 5 min at 1000 rpm at 4°C, the pellet was resuspended in 3 ml of homogenization buffer (0.25 M sucrose, 3 mM imidazole, 1 mM Tris-HCl; pH 7.4, 1 mM EDTA). Cells were homogenized by drawing ∼ 20 times through a 25-gauge needle until the ratio between unbroken cells and free nuclei became 20%:80%. The postnuclear supernatant was obtained by centrifugation at 2,500 rpm at 4°C for 3 min, and then the supernatant was adjusted to 1.4 M sucrose by the addition of ice-cold 2.3 M sucrose in 10 mM Tris-HCl (pH 7.4). Next, 1.2 ml of 2.3 M sucrose at the bottom of the tube was overlaid with 1.2 ml of the supernatant adjusted to 1.4 M sucrose followed by sequential overlay with 1.2 ml of 1.2 M and 0.5 ml of 0.8 M sucrose (10 mM Tris-HCl, pH 7.4). Gradients were centrifuged for 3 h at 182,348 rcf (4°C) in an SW 55 rotor (Beckman Coulter). The turbid band at the 0.8 M/1.2 M sucrose interface containing the Golgi membranes was harvested in ∼500 µl aliquot by syringe puncture.

### In Vitro Furin Cleavage Assay

To investigate whether SP-C is directly cleaved by Furin, EGFP-SP-C was expressed and purified from MLE12 cells by first incubating clarified cell lysates with 15-20 μl GFP-Trap Agarose (ChromoTek) overnight at 4 °C with rotation. Afterwards, beads were recovered by centrifugation, washed three times in washing buffer, and then incubated with 75 μl of Furin cleavage buffer (20 mM HEPES, 1 mM CaCl_2_, 0.2 mM β-mercaptoethanol, 0.1% Triton X-100; pH 7.5) supplemented with 1 U Furin (#P8077S, NEB) for 1 h at 37°C. Furin buffer containing Decanoyl-Arg-Val-Lys-Arg-CMK was included in parallel reactions. The eluted protein in SDS loading buffer with 40 mM DTT was then analyzed via immunoblot.

### Generation of Furin-like proprotein convertase pre-profragments cDNA constructs *and stable cell lines* by lentivirus

pHAGE2-pEF1α-ppPC5-mCherry, pHAGE2-pEF1α-ppPC7-mCherry, and pHAGE2-pEF1α-ppFurin-mCherry were constructed by cloning mouse ppPC5 (1-109 aa), ppPC7 (1-116 aa), and ppFurin (1-142 aa) (IDT gBlock) into the pHAGE2-pEF1α vector. psPAX2 and pMD2G were obtained from Addgene. HEK293T cells were seeded into 10-cm dishes at a seeding density of 300,000 cells/ml on day 1. On day 2, cells were co-transfected with three plasmids, pHAGE2-ppPC5/-7/-Furin-mCherry, psPAX2, and pMD2.G, using JetOptimus (Polyplus-Satorius). Culture medium was changed 24 h post transfection. Lentiviruses were collected after 24 h and 48 h transfection, filtered using a 0.45 µm PES syringe filter, and used for transduction. Proprotein convertase pre-profragment expressing MLE12 cells were generated via transduction using the lentiviral supernatant with 10 µg/ml polybrene, followed by enrichment of mCherry-expressing cells by FACS sorting (CytoFlex SRT) 4 days post-transduction.

### RNA Isolation and Quantitative Real Time Polymerase Chain Reaction

RNA was extracted from cells using RNeasy Mini Kit (Qiagen, Valencia, CA) following the manufacturer’s protocol. The concentration and quality of extracted RNA from lung tissues were measured using NanoDrop® One (Thermo Scientific, Wilmington, DE), and RNA was reverse-transcribed into cDNA using high-capacity (ThermoFisher). RT-qPCR was performed on a QuantStudio 7 Flex Real-Time PCR System in 384 well plate (Applied Biosystems) with results normalized to *Actb* gene expression. Primer sequences for all mouse genes are listed in **Supplemental Table 2.**

### Mouse Alveolar Type 2 Cell Isolation

Flow cytometry was performed as we described (29, 30, 48-50). Blood free perfused lungs were digested in Phosphate Buffered Saline (Mg and Ca free) with 2 mg/ml Collagenase Type I (Gibco Cat# 17100017) and 50 units of DNase (Millipore Sigma Cat# D5025), passed through 70-μm nylon mesh to obtain single-cell suspensions, and then processed with ACK Lysis Buffer (Thermo Fisher). Cell pellets collected by centrifugation were resuspended in PBS and aliquots removed for cell count using a NucleoCounter (New Brunswick Scientific, Edison, NJ). CD45, CD31, CD140α positive cells were removed by negative selection using biotinylated antibodies incubated for 30 mins prior to Dynabead depletion. Fluorophore-conjugated antibodies were used to stain lung cell populations (**Supplemental Table 1)**. All cells were sorted using a CytoFlex SRT (Beckman), and cells were captured in ice-cold FACS buffer (0.1% BSA, 2mM EDTA, and PBS pH 7.4). FACS isolation of distal lung epithelial populations was performed as previously described (51) (pregated on live, singlets, CD45-CD31-Epcam+ cells), AT2 – CD200 ^Hi^, CD104-; AT1 – CD200 ^Lo^, CD104-; ciliated and secretory - CD104+ was performed using a sorting strategy depicted in Supplemental **Figure S1.** qRT-CR analysis was performed on these population to confirm the purity of each group (**Figure S1**).

### Bulk RNA Sequencing (popRNAseq): Sample processing and analysis

A previously published data (JCI AMPK 2025) set was mined for expression of select AT2 specific genes. Briefly, flow sorted AT2 cells (JCI AMPK 2025) were isolated from C57/B6 mice and RNA extracted as described above. Library prep and sequencing was performed by Children’s Hospital of Philadelphia High Throughput Sequencing Core. Resulting fastq files from paired end reads were trimmed, processed and evaluated for quality control using fastp(52). Resulting files were aligned against the mouse genome (mm10) using bowtie2(53) and quantified via featureCount(54). Duplicate reads and chimeric fragments were flagged and excluded from the analysis. Applying limma-voom, read counts were converted to log2-counts-per-million (logCPM) and the mean-variance relationship was modelled with precision weights followed by differential expression analysis (55, 56).

### Statistics

All data are depicted with dot-plots and presented as group mean ± SEM unless otherwise indicated. Statistical analyses were performed with GraphPad Prism (San Diego, CA). Student’s t-test (1 or 2 tailed as appropriate) were used for 2 groups; Multiple comparisons were done by analysis of variance (ANOVA) was performed with post hoc testing as indicated. In all cases statistical significance was considered at p values ≤ 0.05.

### Data and code availability

Analysis of previously published data was performed on GSE296513.

## RESULTS

### Wildtype ProSP-C Is Restricted to Intracellular Organellar Compartments of AT2 Cells

Subcellular localization of proSP-C in AT2 cells in vivo was performed using immunogold electron microscopy of lung tissue from 6–8-week-old SP-C^WT^ and SP-C^I73T^ mice. Using a rabbit polyclonal antibody specific for proSP-C, labeling of wildtype isoforms was readily detected within the Golgi complex multivesicular bodies (MVBs), and lamellar bodies (LB) (**Figure 1A-B, panels i, ii, and iii**). Notably, specific labeling for wild-type (WT) SP-C was not detected over nuclei, mitochondria, or at the plasma membrane (**Figure 1A, panel iii**) supporting the concept of direct anterograde trafficking to LB under homeostatic conditions. In contrast, in addition to Golgi and MVB, the I73T mutant SP-C localized to apical tubular structures beneath the plasma membrane, extending along the microvilli of the AT2 cell surface **(Figure 1B, panel iii).**

**Figure 1.**
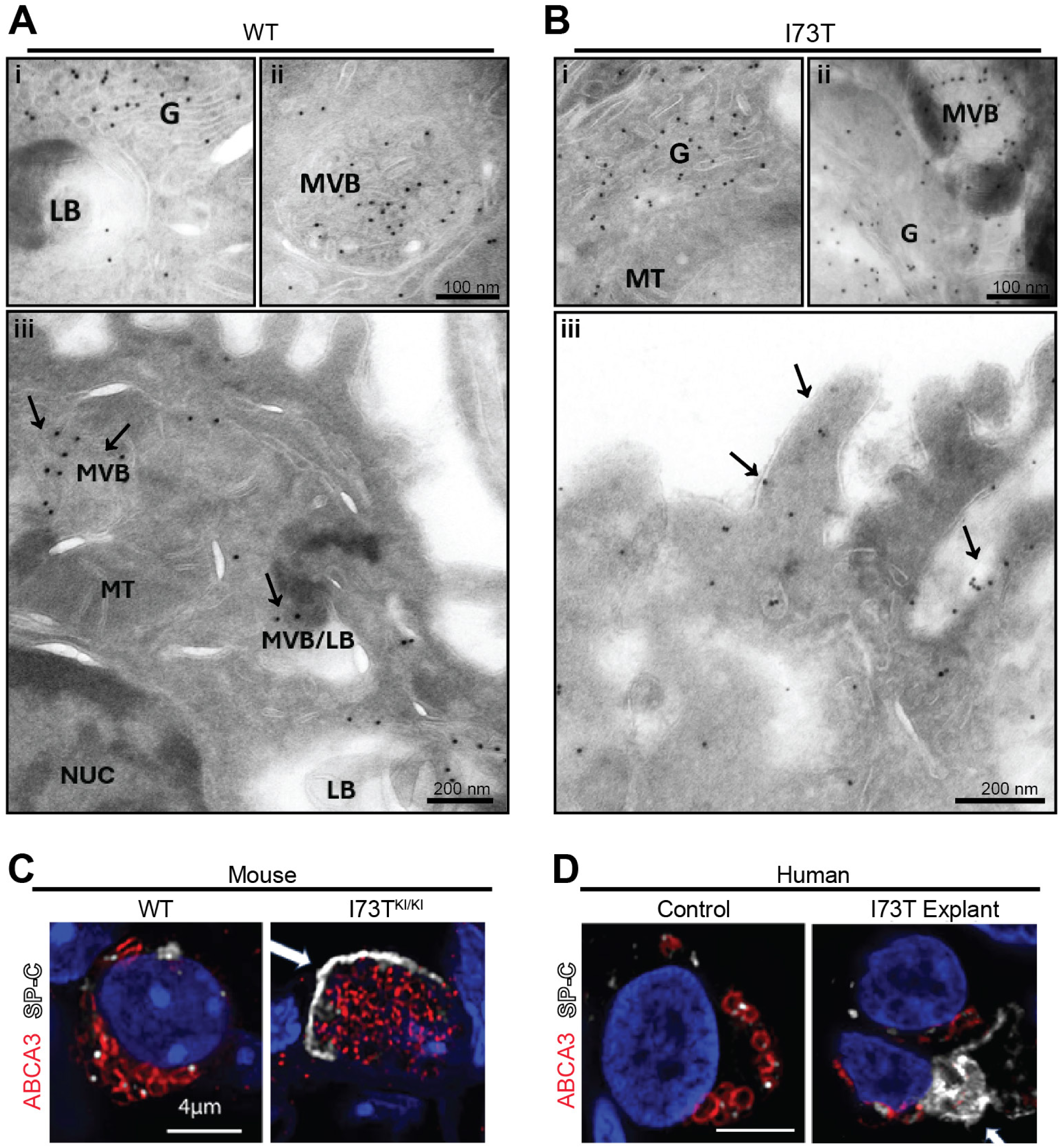
Immunogold localization of Pro-SP-C in AT2 cells. **[A]** Lungs of 6-8 week old SP-C^wt/wt^ mice and **[B]** SP-C^I73T^ ^KI/KI^ mice were fixed, ultra-thin sectioned and labelled using rabbit polyclonal antibodies directed against NH_2_-terminus of Pro-SP-C and protein and protein A gold (10nm). Wildtype SP-C was absent from the cell surface. LB: lamellar body; MVB: multivesicular body; G: golgi complex; NUC: nucleus; MT: mitochondria. Scale bar is 0.2 µm. Representative images of AT2 cells stained for lamellar body marker ABCA3 (red) and SP-C (white) in tissues sections from [C] wildtype and I73T mouse lung and **[D]** human I73T explant lung. Nuclei are counterstained with DAPI (blue). Scale bar: 4µm.

To complement this, confocal immunofluorescence microscopy was performed on tissue sections from both WT and Sftpc^I73T^ mice and human lung specimens. In wild-type mouse AT2 cells, proSP-C staining was found within ABCA3-positive LB (**Figure 1C**). In contrast, sections from SftpcI73T KI/KI mutant mice displayed aberrant accumulation of proSP-C near the cell periphery (**Figure 1C**, arrow). A similar mislocalization pattern was observed in sections from a human lung explant carrying the I73T *SFTPC* mutation, but not in a normal donor (*SFTPC* WT/WT) lung (**Figure 1D**). Together, these findings demonstrate that while proSP-C is normally restricted to intracellular organelles, the pathogenic I73T variant disrupts normal trafficking, promoting plasma membrane retention in both mouse and human AT2 cells.

### Trafficking of SFTPC Isoforms in MLE-12 Cell Lines

To investigate the maturation, processing, and trafficking of *SFTPC* variants, we generated doxycycline-inducible MLE-12 cell lines expressing previously published GFP-tagged human WT and two mutant (I73T and C121G) *SFTPC* constructs (22) (**Figure 2A**). Similar mRNA expression levels of *SFTPC* were observed across the cell lines post-24h doxycycline induction (**Supplemental Figure S2**). Western blotting revealed that GFP-SP-C^WT^ is properly processed, displaying a banding pattern consistent with the expected primary translation product (∼48 kDa), a palmitoylated pro-form, and two processing intermediates (**Figure 2B**) (57). In contrast, expression of the GFP-SP-C^I73T^ variant generated higher molecular weight pro-protein forms, including a prominent band previously shown to be glycosylated (35), all indicative of altered protein maturation. The GFP-SP-C^C121G^ mutant expressed as a single primary translation product consistent with ER retention and lack of further processing including Golgi palmitoylation (22, 25, 30, 50).

**Figure 2.**
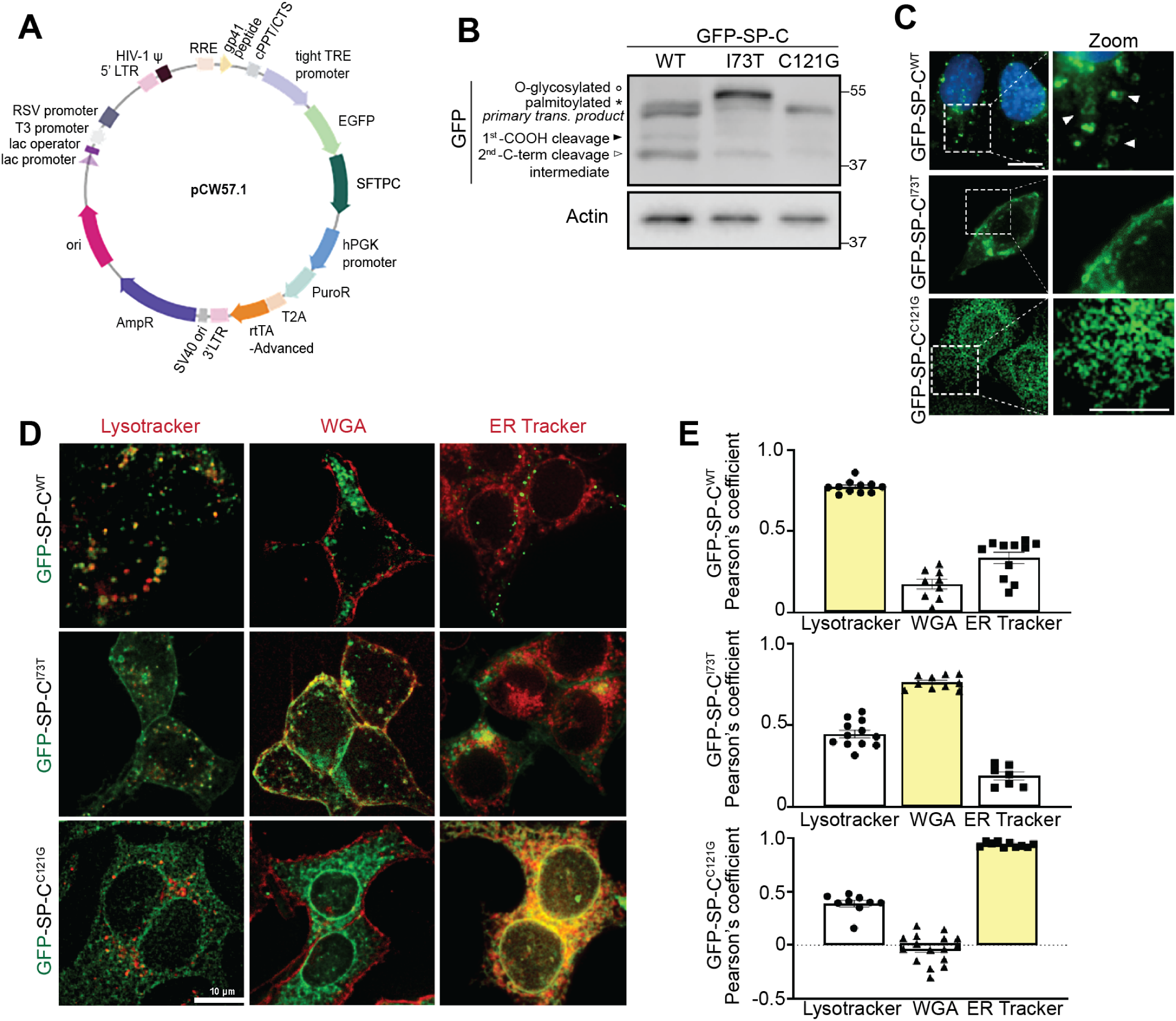
Generation and characterization of SFTPC-expressing MLE-12 cell lines. **[A]** Schematic of GFP-tagged Surfactant Protein C constructs cloned into the lentiviral expression vector pCW57.1 with a Tet-responsive element (TRE). **[B]** Immunoblot of GFP-tagged SP-C WT, I73T, and C121G constructs expressed in MLE-12 cells for 24h with 2.5µM doxycycline induction. Arrowheads and asterisk denote C-terminal processing intermediates and the palmitoylated pro-form, respectively. **[C]** Live-cell fluorescent widefield microscopy of GFP-SP-C localization in MLE-12 cells. GFP signal (green) and Hoechst-stained nuclei (blue) are shown. White arrow heads denote wildtype localization of SP-C on membranes of vesicles. Scale bar: 10 μm (top), 5 μm (bottom inset). **[D]** Co-localization of GFP-SP-C WT, I73T, and C121G isoforms following 24h of doxycycline induction in live cells co-stained with organelle markers: LysoTracker (acidic LROs), Wheat Germ Agglutinin (WGA) (plasma membrane), and ER Tracker (endoplasmic reticulum). Merged images show organelle marker (red) and GFP-SP-C (green). Scale bar: 10 μm. **[E]** Quantification of co-localization by Pearson’s correlation coefficient between GFP signal and each organelle marker in WT, I73T, and C121G-expressing cells. Data represent mean ± SEM from *n* ≥ 10 cells per condition, pooled from 3 independent experiments.

Live-cell immunofluorescence microscopy of MLE-12 cells (**Figure 2C**) demonstrated distinct subcellular localization patterns for each of the variants. GFP-SP-C^WT^ predominantly localized to ring-shaped peripheral vesicular structures while the GFP-SP-C^I73T^ mutant displayed abnormal accumulation on the plasma membrane, and the C121G mutant was proximally retained within the endoplasmic reticulum (ER). Quantitative co-localization studies using WGA, Lysotracker, and ER-tracker (**Figure 2D-E**) confirmed these patterns with EGFP-SFTPC^WT^ localizing to lysosome-related organelles (LROs), EGFP-SFTPC^I73T^ retained on WGA (+) plasma membranes, and EGFP-SFTPC^C121G^ confined to the ER.

### Pitstop2 Selectively Alters Trafficking of the SFTPC^I73T^ Mutant Isoform

To further characterize the observed accumulation of SP-C^I73T^ on the plasma membrane and to exclude the plasma membrane as a transient routing site for SP-C^WT^, we inhibited clathrin-mediated endocytosis using Pitstop 2 (58, 59). We first validated the efficacy of Pitstop 2 in MLE-12 cells by assessing endocytic uptake of a non-SP-C related cargo. Cells were treated with 20µM Pitstop 2, then incubated with AlexaFluor488-conjugated mouse transferrin for 30 minutes, washed, and fixed for imaging. Pitstop 2–treated cells showed markedly reduced transferrin uptake compared to vehicle-treated controls, confirming effective inhibition of endocytosis (**Supplemental Figure S3**).

GFP-SP-C expression was then induced in MLE12 lines using doxycycline overnight prior to Pitstop 2 treatment for 30 minutes. Immunofluorescence analysis (**Figure 3A**) showed that Pitstop2 further enhanced deposition of GFP-tagged I73T mutant protein in a contiguous plasma membrane pattern, corroborated by increased surface GFP intensity (**Figure 3C**), while total GFP-SP-C levels remained unchanged (**Figure 3B**), indicating no effect on protein synthesis or degradation during the short treatment period. In contrast, Pitstop 2 treatment had no effect on the intracellular vesicular distribution pattern or signal intensity of GFP-SP-C^WT^. Similarly, Pitstop 2 did not alter the ER retention of GFP-SPC^C121G^ (not shown) supporting the selective plasma membrane localization of the I73T variant.

**Figure 3.**
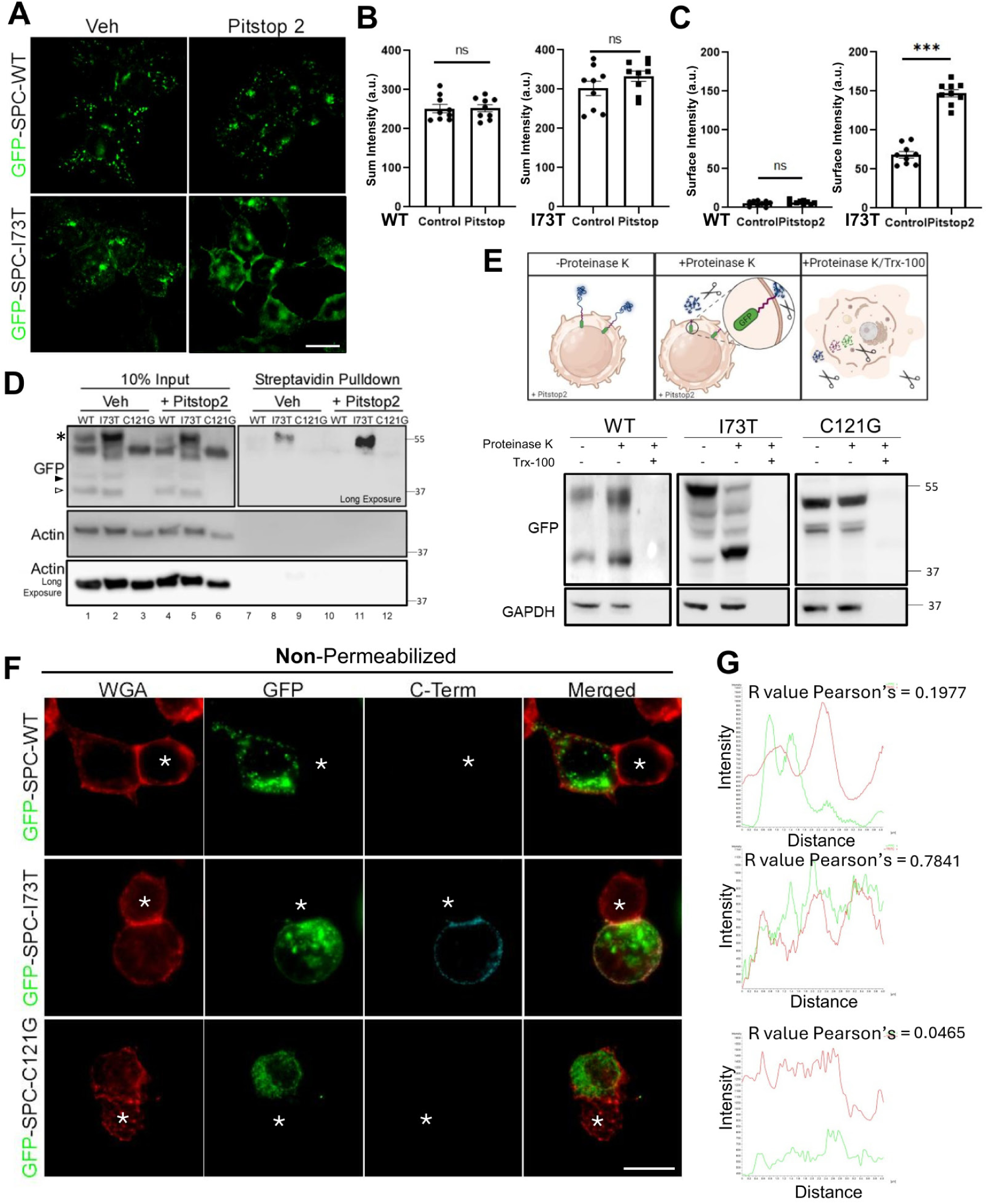
Divergence of wildtype and mutant Pro-SP-C expression patterns. **[A]** 24h post-doxycycline induction of GFP-SP-C^WT^ and GFP-SP-C^I73T^ in MLE-12s, cells were treated with vehicle (DMSO) or Pitstop 2 for 2h. **[B]** Quantification of total GFP signal and **[C]** contiguous plasma membrane GFP signal for WT and I73T. **[D]** MLE-12 cells were treated with doxycycline for 24 h to induce SP-C expression, followed by a 2 h co-treatment with Pitstop 2 or vehicle. Surface proteins were biotinylated, enriched via streptavidin pulldown, and immunoblotted for GFP-tagged SP-C. **[E]** GFP-SP-C at the cell surface was assessed via susceptibility to proteolysis by Proteinase K protection assay in non-permeabilized vs. Triton X-100– permeabilized cells. **[F]** Immunofluorescence analysis in non-permeabilized MLE-12 cells expressing GFP-SP-C variants. WGA (red) marks plasma membrane. Anti-GFP (green) detects N-terminus of SP-C; anti–C-terminal antibody against SP-C (cyan). Asterisks mark non-transfected cells. Yellow arrows denote orientation of line scans at the plasma membrane **[G]** Intensity profiles of line-scan analyses across cell membranes in non-permeabilized cells. Pearson’s correlation coefficients (R values) between GFP (green) and WGA (red) channels are shown for each variant. All quantitation data are shown as mean ± SEM from three independent experiments. Statistical analyses are performed using two-tailed Student’s t-test. *, p < 0.05; **, p < 0.01; ***, p < 0.001; n.s., not significant.

### The SFTPC^I73T^ Mutant Isoform Is Restricted to the Plasma Membrane

To further characterize SP-C^I73T^ mislocalization, we performed cell surface biotinylation followed by streptavidin pull-down assays. Immunoblotting for GFP demonstrated that only SP-C^I73T^ isoforms reach the cell surface which was amplified by PitStop 2 treatment (**Figure 3D**). Further assessment of cell surface deposition was done using a protease protection assay performed by treatment of SP-C MLE-12 cells with proteinase K (**Figure 3E**), which resulted in a prominent band shift of the proSP-C^I73T^, indicating its presence on the plasma membrane and extracellular exposure. In contrast, both GFP-SP-C^WT^ and the ER retained GFP-SP-C^C121G^ mutant were resistant to proteinase K cleavage, demonstrating intracellular restriction and protection.

The cell surface orientation of mutant SFTPC^I73T^ was obtained using non-permeabilized cells maintained at 4°C and co-stained with an epitope-specific proSP-C antibody recognizing the COOH-terminus (40), with labeling by WGA marking the cell surface (**Figure 3F-G**). In contrast to SP-C^WT^ or SP-C^C121G^, quantitatively significant COOH-terminus proSP-C staining co-localizing with WGA was only observed in GFP-SP-C^I73T^ mutant-expressing MLE-12 cells, indicating that the cell surface-localized isoform retains its BRICHOS domain which is accessible to the extracellular space.

### Trafficking Divergence of SFTPC Isoforms Is Recapitulated in iPSC-derived human AT2 cells

The trafficking and processing of SFTPC isoforms was next studied in a published translationally relevant in vitro model, iPSC-derived alveolar type 2 (iAT2) cells expressing either tdTomato/WT or I73T/tdTomato SFTPC (23). Immunofluorescence staining for proSP-C and E-cadherin (**Figure 4A**) revealed that in tdTomato/WT iAT2 cells, proSP-C expression was again restricted to intracellular vesicular compartments. Treatment with the lysosomal v-ATPase inhibitor Bafilomycin A1 (BafA1), which de-acidifies lysosomes thereby inactivating acid-dependent lysosomal hydrolases found in LB such as Cathepsins (33) (60), caused an accumulation of proSP-C within enlarged, swollen vesicles. In contrast, and consistent with our previous findings in MLE-12 cell lines, Pitstop 2 treatment did not significantly change the localization of wild-type SP-C but again specifically altered the cellular distribution of the SP-C^I73T^ mutant by enhancing its cell-surface expression.

**Figure 4.**
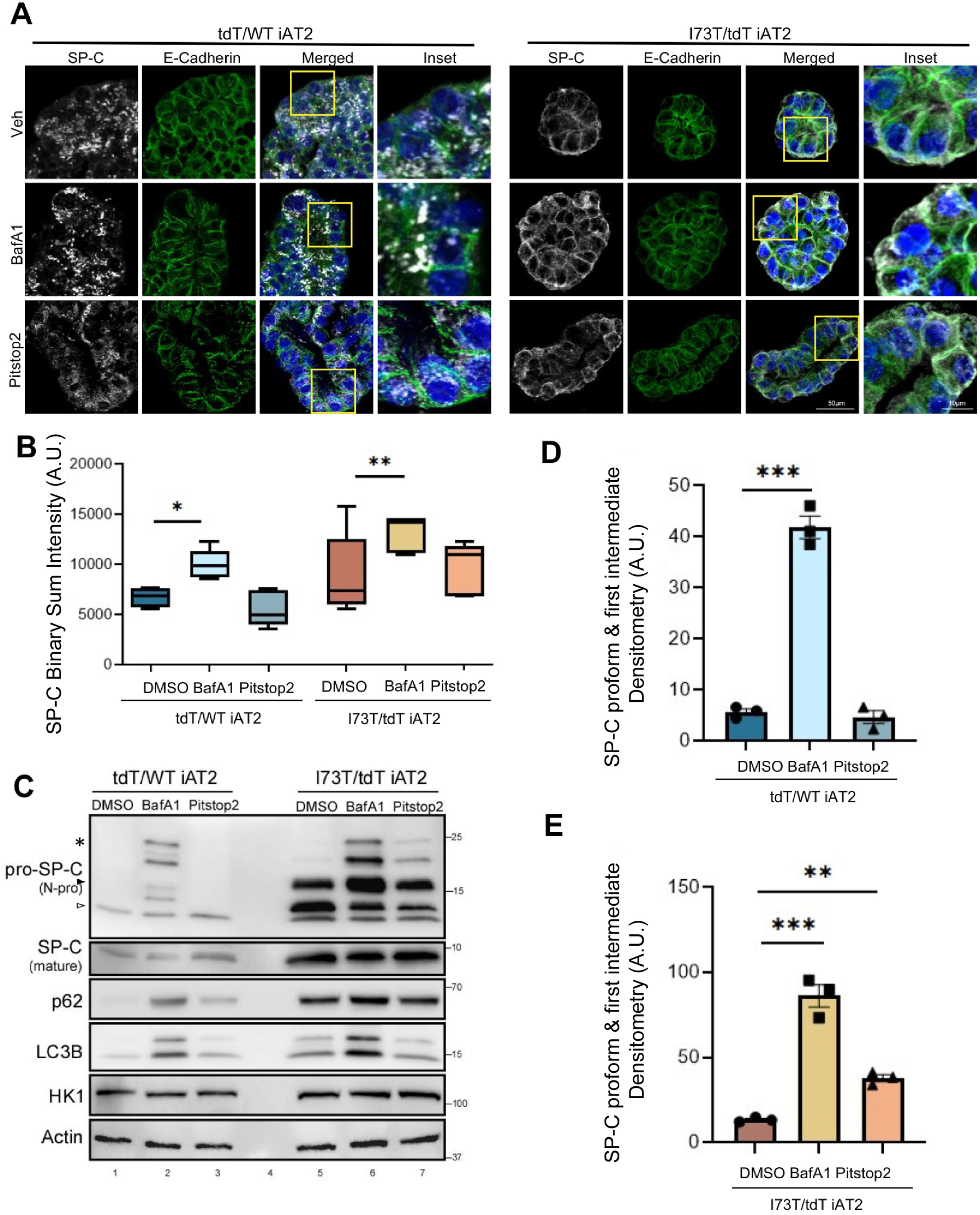
Inhibition of lysosomal acidification and endocytosis differentially alters SP-C processing in WT and I73T iAT2 cells. **[A]** Confocal microscopy of WT and I73T iAT2 alveolospheres treated with DMSO, Bafilomycin A1 (BafA1) (50nM) 16h, or Pitstop 2 (20µM) for 2h. E-cadherin (green), nuclei (Hoechst, blue), and SP-C (N-Pro, white). Scale bars: 50µm, 10 µm for inset. **[B]** Quantification of SP-C puncta intensity across conditions, shown as binary sum intensity per alveolosphere area from 3 different replicates. *P < 0.05, **P < 0.01 [one-way ANOVA with Fisher’s LSD post-hoc test]. **[C]** Immunoblot of SP-C processing intermediates (*= palmitoylated pro-form, arrowheads= C-term cleavage intermediates) and mature SP-C in WT and I73T iAT2s after treatment. Autophagy markers p62 and LC3B, HK1 and Actin were included as controls. **[D-E]** Densitometric analysis of pro-form and first-intermediate SP-C bands from (C; *brackets*) normalized to Actin. Data represent mean ± SEM from *n* = 3 experiments. Statistical comparisons were performed using one-way ANOVA and Sidak post hoc correction and significance denoted as *, p < 0.05; **, p < 0.01; ***, p < 0.001 versus control.

Quantification of SP-C fluorescence intensity (**Figure 4B**) coupled with assessment of proSP-C isoforms by Western blotting (**Figure 4C-E**) confirmed trafficking and maturation differences. In the tdTomato/WT iAT2 cells, both pro-SP-C and mature SP-C forms were detected, with a notable increase in pro-SP-C isoforms (*bracket;* } highlights accumulated SP-C pro-form & intermediate) and a decrease in mature SP-C following BafA1 treatment consistent with inhibition of late NH2 propeptide processing events **(Figure 4C**). At baseline SFTPC^I73T^-iAT2 cells accumulated significant amounts of proSP-C^I73T^ isoforms consistent with aberrant processing, which were increased by BafA1, suggesting some of these processed forms reside in acidic compartments. Consistent with previously published data (23, 28), SP-C^I73T^-iAT2 cells also demonstrate higher degrees of autophagic flux as evidenced by greater BafA1-induced increases in LC3B and p62 compared with isogenic corrected SFTPC^WT^-iAT2 cells (**Figure 4C**). While Pitstop2 treatment selectively altered proSP-C^I73T^ localization (**Figure 4A**), autophagic markers p62 and LC3B remained unchanged in both *SFTPC* genotypes (**Figure 4C**), indicating that the accumulation of proSP-C^I73T^ under Pitstop2 was specific to SP-C processing and trafficking rather than general autophagy modulation. Together, these results suggest that evoked LB (LRO) dysfunction broadly impairs both proSP-C^WT^ and proSP-C^I73T^ maturation, while inhibition of endocytic trafficking uniquely drives aberrant surface accumulation of proSP-C^I73T^ mutant.

### Post-translational Processing of SP-C involves initial cleavage of the COOH propeptide in the Trans-Golgi Network (TGN)

To pinpoint subcellular sites of potential SP-C COOH-propeptide cleavage events, we evaluated the localization and processing of proSP-C in GFP-SP-C^WT^ MLE-12 cells subjected to temperature shifts and pharmacological perturbations. Immunofluorescence analysis (**Figure 5A**) showed that treatment with Brefeldin A, which collapses the cis- and medial-Golgi into the ER, resulted in retention of proSP-C^WT^ within the ER. Meanwhile incubation at 20°C, a condition shown to block protein transport beyond trans-Golgi/TGN (61-64), led to pronounced enrichment of proSP-C^WT^ within the Golgi/TGN.

**Figure 5.**
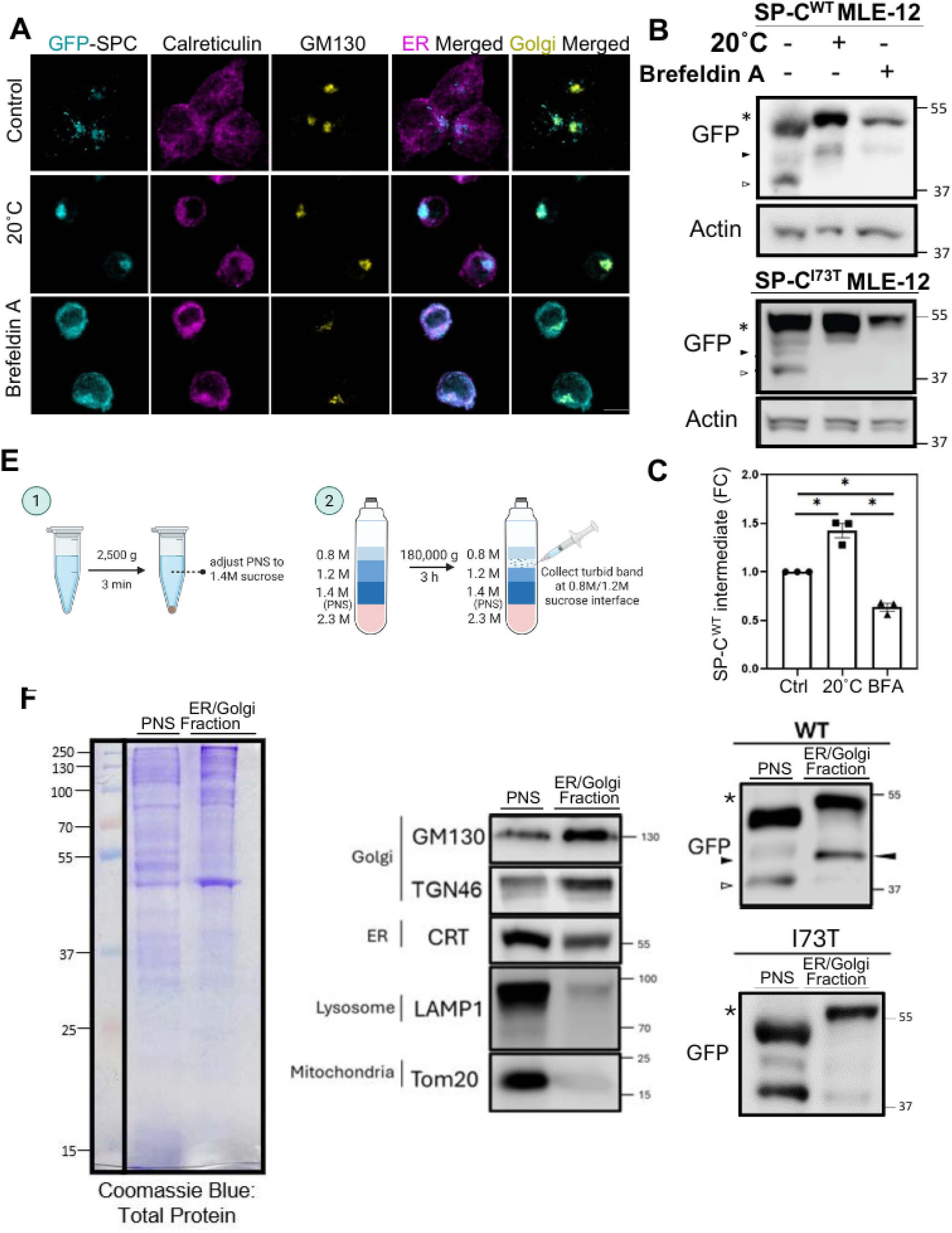
First cleavage of the SP-C COOH-propeptide occurs in the trans-Golgi network. **[A]** Representative images of MLE-12 cells induced with doxycycline for 3 h followed by trans-Golgi trafficking inhibition at 20°C for 2h, or after cis-/medial-Golgi collapse induced by Brefeldin A for 2h, showing GFP-SP-C (cyan) localization relative to the ER (Calreticulin, magenta) and Golgi (GM130, yellow) as compared to control condition (37°C). Scale bar: 10 μm. **[B]** Representative immunoblot of GFP-SP-C wildtype and I73T mutant isoform following treatment (A) and **[C]** densitometry analysis of wildtype SP-C first COOH-cleavage intermediate band (solid arrowhead) normalized to actin, shown as fold change relative to control. Experiments were performed in triplicate and statistical analyses were performed using one-way ANOVA. *, p < 0.05. **[D]** Schematic of sucrose gradient-based subcellular fractionation to isolate ER/Golgi-enriched membranes from postnuclear supernatants (PNS). **[E]** Left: Coomassie staining of total protein in input and ER/Golgi fractions. Middle: Immunoblots showing enrichment of ER, Golgi, and absence of markers from other organelles (mitochondria and lysosomes) in purified fractions. Right: Representative immunoblot showing enrichment of first COOH-cleavage SP-C intermediate (solid arrowhead) in the ER/Golgi fraction of wildtype SP-C expressing MLE-12, and in contrast sparsely detected from this fraction for the I73T mutant.

Western blot analysis with quantitation (**Figure 5B-C**) indicated that GFP-tagged SP-C^WT^ levels increased in cells subjected to the 20°C condition, manifested as elevated levels of both the primary translation product and an intermediate bearing a partially cleaved COOH-propeptide (closed arrowhead) coupled with absence of a more extensively COOH cleaved isoform (open arrowhead) implicating the late Golgi/TGN as the site of initial proSP-C processing. Meanwhile, a 20°C temperature block failed to enrich the partially cleaved C-terminal propeptide intermediate of mutant SP-C^I73T^, indicating that this variant deviates from the normal maturation pathway of WT SP-C at an early post-ER stage. Notably Brefeldin A treatment further inhibited proSP-C remodeling preventing enrichment of the cleaved band further supporting the requirement for delivery to trans-Golgi/TGN to effect COOH-propeptide cleavage.

To further validate these findings, we next performed subcellular fractionation of GFP-SP-C^WT^ and GFP-SP-C^I73T^ expressing MLE-12 cells (**Figure 5D**). Purity of the fractions, assessed using immunoblotting for organelle specific markers, confirmed the identity of the ER/Golgi, evidenced as enrichment in GM130, TGN46, and Calreticulin with minimal contamination by lysosomes (LAMP1) or mitochondria (Tom20), identified in the unfractionated post-nuclear supernatant (PNS) (**Figure 5E, center panel**). Interestingly compared to PNS, ER / Golgi fractions from GFP-SP-C^WT^ cells were highly enriched in both a full length proSP-C isoform and an intermediate bearing a partially cleaved COOH-propeptide (closed arrowhead) but with absence of the lowest MW isoform. In contrast, no proSP-C processing intermediates were identified in ER/Golgi fractions from GFP-SP-C^I73T^ cells. Together, these results demonstrate biochemically that the initial cleavage of the COOH-propeptide of SP-C occurs within a proximal compartment (Golgi/TGN) but that mutant proSP-C^I73T^ is excluded from this initial processing in this compartment.

### Mutant I73T SP-C expression disrupts Golgi morphology

To assess whether expression of the SP-C^I73T^ mutant alters Golgi structure, we examined the ultrastructure of murine AT2 cells by transmission electron microscopy (TEM) from WT and I73T *SFTPC* mice. In wildtype mice, the Golgi apparatus appeared as organized stacks of flattened cisternae with closely apposed membranes (**Figure 6A, top panels**). In contrast, SP-C^I73T^-expressing AT2 cells exhibited a fragmented Golgi morphology, characterized by dispersed and swollen cisternae with loss of the regular stacked architecture (**Figure 6A, bottom panels**).

**Figure 6.**
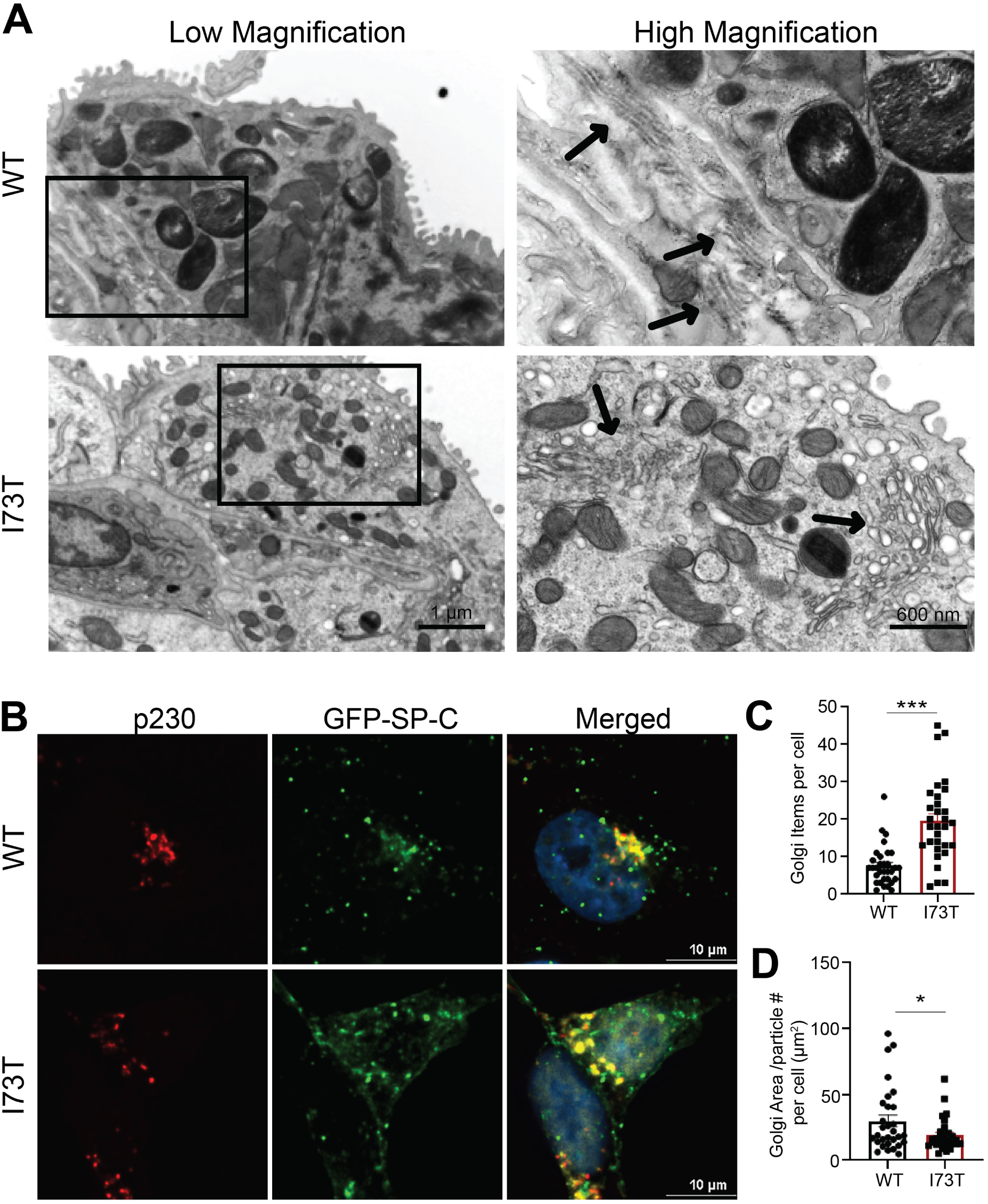
I73T SP-C expression disrupts Golgi architecture in MLE-12 cells. **[A]** Representative transmission electron microscopy (TEM) images of murine AT2 cells from whole lung mounts. Arrows point to Golgi. **[B]** Confocal images of MLE-12 cells expressing GFP-tagged WT or I73T SP-C (green) and immunofluorescence stained for the trans-Golgi marker p230 (red). Nuclei are stained with Hoechst (blue). [**C, D]** Quantification of the number of p230-labeled Golgi elements per cell (C) and Golgi area/ number of Golgi particles within a cell (D). All quantitation data are shown as mean ± SEM from three independent experiments. Statistical analyses are performed using two-tailed Student’s t-test. *, p < 0.05; ***, p < 0.001.

We next visualized Golgi organization using immunofluorescence staining for the trans-Golgi marker p230. In WT SP-C–expressing MLE-12 cells, p230 staining was concentrated in a compact perinuclear ribbon that partially colocalized with GFP-SP-C (**Figure 6B**). In SP-C^I73T^ -expressing cells, the p230 signal was more dispersed, with smaller and more numerous Golgi elements distributed throughout the cytoplasm. Quantitative analysis confirmed a significant increase in the number of discrete Golgi puncta per cell in SP-C^I73T^ -expressing cells compared to WT (**Figure 6C**) and a reduction in average Golgi area per particle (**Figure 6D**), consistent with Golgi fragmentation. Given the Golgi’s central role in directing cargo to downstream compartments, we extend prior reports of endolysosomal perturbations (27, 28) by showing that these defects are accompanied by Golgi fragmentation.

### Furin Proprotein Convertase Participates in proSP-C Processing

Having observed that SP-C^I73T^ expression and alterations in its post-translational processing are associated with disruptions in Golgi morphology, we next focused on defining the early proteolytic processing events that occur in this compartment to provide insight into SP-C maturation under physiologic conditions. SP-C is a member of the BRICHOS family of proteins, several of which are known to undergo cleavage by furin-like proprotein convertases (PPCs) (65). These calcium-dependent serine endoproteases, which function predominantly in the trans-Golgi network (TGN), recognize multibasic motifs and play a central role in activating many secretory and transmembrane proteins within the distal secretory pathway(66). Based on this, we investigated the involvement of furin-like proprotein convertases (PCs) in the initial cleavage and processing of SP-C using both pharmacological and genetic approaches. Treatment of MLE-12 cells expressing GFP-tagged WT and I73T SP-C with the pan-proprotein convertase inhibitor (DC1)(67) resulted in a loss of cleaved proSP-C isoforms indicating that furin-like PPC activity is required for SP-C processing (**Figure 7A-B**). Similar results were obtained using primary murine AT2 (mAT2) cells treated with DC1 (**Figure 7C-D**). Using the MLE-12 GFP-SP-C^WT^ cell line, treatment with DC1 at the time of doxycycline induction resulted in accumulation of GFP-SP-C^WT^ at the Golgi after 9 hours, as indicated by colocalization with the cis-Golgi marker GM130 (**Figure 7E**) compatible with delayed exit of SP-C from the Golgi compartment indicating that inhibition of proprotein convertases (PPCs) impacts both the processing and trafficking kinetics of SP-C.

**Figure 7.**
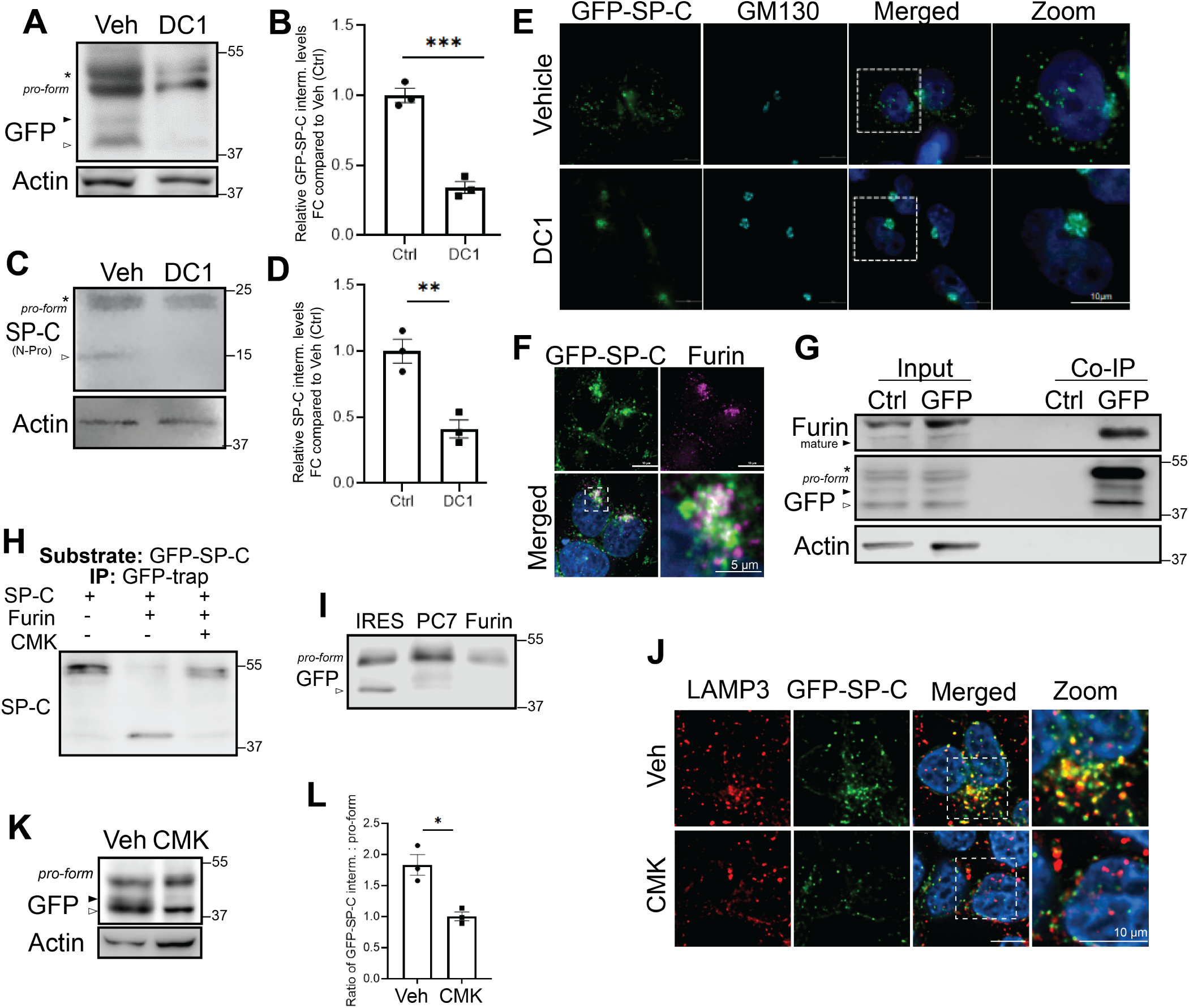
Furin-like pro-protein convertases are candidate enzymes for initial SP-C cleavage. **[A-B]** Western blot analysis and quantitation of c-term SP-C cleavage (arrowheads) in MLE-12 cells co-treated with DC1 (dicoumarol-related pan-proprotein convertase inhibitor) and doxycycline to induce the expression of GFP-SPC^WT^ for 9 h. **[C-D]** Immunoblot and quantitation of c-term SP-C cleavage after overnight DC1 treatment in primary murine AT2 isolated from SP-C^WT^ mice shows similar inhibition of SP-C processing. **[E] I**mmunofluorescence of GFP-SPC^WT^ MLE-12 cells co-treated with Doxycycline and DC1 for 9h, fixed and stained with GM130 (Golgi marker). **[F]** Immunofluorescence of GFP-SPC^WT^ after 6h of doxycycline-induction (green) shows colocalization with Furin (magenta) in the perinuclear region of MLE12 cells. Hoechst stains nuclei. Scale bars: 10 μm **[G]** GFP-pulldown in MLE12 cells using GFP-nanotrap or Control Binding-nanotrap beads shows interaction of mature furin with GFP-tagged SP-C. **[H]** Furin cleavage assay performed with SP-C pro-translation product as the *in-vitro* substrate in the presence and absence of Decanoyl-RVKR-CMK. **[I]** Immunoblot after GFP-pulldown enrichment following 9h of doxycycline showing SP-C processing in MLE-12 cells transduced with individual pre-profragment of PC7 or Furin or IRES empty vector control. **[J]** Immunofluorescence of GFP-SPC^WT^ after overnight of doxycycline-induction and Decanoyl-RVKR-CMK Furin inhibitor stained for LAMP3. **[K]** Immunoblot of GFP-SPC^WT^ expressing MLE-12 cells after overnight treatment of doxycycline and Decanoyl-RVKR-CMK. **[L]** Quantification ratio of SP-C cleaved intermediate relative to pro-form SP-C in each condition.

Transcriptomic profiling using both reanalysis of our published popRNASeq dataset [GSE296513] and qPCR of lung epithelial cells (**Supplemental Figure S4A-C**) confirmed that two proprotein convertases, Furin (*Pcsk3*) and PC7(*Pcsk7*), are each expressed in both mAT2 and MLE-12 cells at significantly higher levels than other PPCs, with Furin showing the highest expression. A transcriptomic survey of human AT2 cells confirms that Furin (*Pcsk3*) is also the highest expressed family member (**Supplemental Figure S4D**). Confocal imaging (**Figure 7F**) showed partial co-localization of GFP-SP-C^WT^ with Furin in a perinuclear region, consistent with Golgi/TGN localization. Co-immunoprecipitation assays confirmed a direct interaction between GFP-SP-C^WT^ and mature Furin (**Figure 7G**). Finally, in vitro cleavage assays demonstrated that recombinant Furin efficiently processes full-length GFP–proSP-C, and this activity was specifically inhibited by the Furin inhibitor (CMK) (**Figure 7H**).

To dissect the contribution of Furin to proSP-C processing, we generated expression constructs encoding for dominant-negative pre-prosegments of Furin and PC7, (**Supplemental Figure S4E**) which act as enzyme specific competitive PC inhibitors (68). RT-PCR confirmed robust expression of each pre-profragment in transfected MLE12 cells (**Supplemental Figure S4F**). When expressed in Dox-inducible SP-C MLE-12 cells Furin pre-prosegment robustly reduced proSP-C cleavage compared to IRES controls (**Figure 7I**). When the pre-prosegments (pp) of Furin was expressed along with ppPC7 and ppPC5, no cleavage of SP-C was detected at early time-points (9 hrs post-dox induction), suggesting functional redundancy among convertase family members (**Supplemental Figure S4G**). At steady state, prolonged inhibition of Furin activity with CMK resulted in accumulation of the full-length proprotein and a concomitant reduction in the second processing intermediate compared to controls (**Figure 7K, L**). Immunofluorescence analysis further revealed decreased colocalization of proSP-C with LAMP3⁺ compartments following Furin inhibition, consistent with impaired progression to distal secretory and lysosome-related organelles (**Figure 7J**). When combined with our earlier time-course data showing delayed Golgi exit upon acute inhibition, these findings indicate that loss of Furin activity not only alters the efficiency of initial C-terminal cleavage but also impacts the kinetics and subcellular distribution of SP-C. Together, these data demonstrate that Furin, a proprotein convertase localized to the Golgi mediates the initial proteolytic processing of SP-C in the late Golgi/TGN.

### Localization of COOH Propeptide Cleavage to the SFTPC BRICHOS Domain

To identify the initial proSP-C COOH-terminal cleavage site for human proSP-C, we first performed an *in silico* prediction using DeepPeptide, which indicated a likely cleaved region between amino acids 150–197 (**Supplemental Figure S5A**). To further refine this prediction, ProP 1.0 algorithim, an artificial neural network program (69) was used to predict potential proprotein convertase cleavage sites within the BRICHOS domain containing the cleavage motif (K/R)-(X)n-(K/R)↓ (where n is 0, 1, 2, 4 or 6 and X is any amino acid). The analysis highlighted a potential cleavage site at residue 167 with the highest score (**Supplemental Figure S5B**). A multiple sequence alignment across species revealed strong conservation of basic residues within this region of the predicted cleavage motif (**Supplemental Figure S5C**). WebLogo analysis visualizing residue frequency confirmed a highly conserved basic stretch consistent with furin-like proprotein convertase recognition (**Supplemental Figure S5D**).

To experimentally test the functional relevance of this predicted site, we next mutated conserved lysine 160 and arginine 167 to alanine in our EGFP-SFTPC-tagged SP-C constructs. Western blot analysis of lysates prepared following transient transfection of MLE-12 cells demonstrate significant accumulation of the unprocessed palmitoylated EGFP-proSFTPC^R167A/K160A^ mutant (**Supplemental Figure S5E**) indicating physiological cleavage of native proSP-C likely occurs at position 160/167 within the COOH BRICHOS domain and represents a critical early processing step in SP-C maturation.

## DISCUSSION

In this study, we investigated the biosynthetic steps that govern SP-C maturation and the impact of disease related SP-C mutants on these events with a focus on understanding the post-translational trafficking itinerary and early proteolytic processing of the COOH-terminal propeptide. In doing so we were able to define key differences in trafficking and post-translational processing between WT SP-C and a well described interstitial lung disease-associated *SFTPC* mutant, SP-C^I73T^. In 3 separate models including primary murine and human iPSC-derived AT2 cells, SP-C^WT^ was highly concentrated in acidic LROs while the SP-C^I73T^ isoform accumulated on the plasma membrane **(Figures 2 and 4)**. This observation was then corroborated by 5 separate biochemical and cell biological assays including inhibition of clathrin-mediated endocytosis, surface biotinylation, immunogold EM, immunofluorescent staining of non-permeabilized cells, and proteinase K protection assays, collectively supporting divergence of SP-C^I73T^ trafficking from SP-C^WT^ (**Figures 3 and 4**). Then, utilizing Brefeldin A, temperature shifts, and subcellular fractionation of doxycycline-inducible mouse lung epithelial (MLE-12) cell lines expressing either WT SP-C or SP-C^I*73T*^, we determined that exclusion of proSP-C^I73T^ from normal anterograde routing occurs very early in the biosynthetic pathway prior to any proteolytic proprotein processing which begins in the Golgi (**Figure 5**). Finally, we hypothesized and confirmed using functional assays that the early propeptide COOH cleavage event involves Furin (*Pcsk3*), the most highly enriched member of the proprotein convertase family in AT2 cells (**Figure 6**). Collectively, our data demonstrate that trafficking pathways for maturation of WT and mutant SP-C^I*73T*^ diverge prior to the TGN, where initial cleavage of the COOH-terminal SP-C propeptide occurs.

The localization of SP-C^I*73T*^ to the cell surface appears to be specific to the mutant I73T isoform. In contrast to a recent report (35) under any conditions tested here across 3 lung epithelial models, we failed to detect significant amounts of SP-C^WT^ reaching the plasma membrane, and our data corroborate multiple prior reports documenting classic anterograde trafficking routes for proSP-C from Golgi via MVB to LB (9, 22, 70). Potential reasons for this discrepancy likely reside in the unique structure and biology of SP-C and the model systems chosen. In particular, the primary experimental model used by Dickens, et al (35) was based on exogenous expression of SP-C in HeLa cells that do not contain LBs, a lysosome-related organelle (LRO) found in AT2 cells. Emerging evidence suggests that the fidelity of LB cargo trafficking depends on specialized anterograde pathways that are only preserved in models containing LROs (71). Thus, accurate recapitulation of SP-C trafficking requires cell systems that retain the organelle architecture and sorting machinery of the alveolar epithelium.

In addition, we chose to enhance visualization of any potential transient interaction with proSP-C at the plasma membrane through inhibition of endocytosis using Pitstop 2 (58, 59). This small-molecule strategy to block clathrin-mediated endocytosis offers an advantage over genetic approaches such as CRISPR knockouts or knock-sideways depletion of adaptor proteins like AP-2, which can cause broader trafficking defects that disrupt normal SP-C maturation, including that of key sorting receptors such as LIMP2 (for lysosomal hydrolases) and LAMPs (72). Thus, transient endocytosis inhibition offers a more precise method to interrogate these trafficking defects without disrupting global sorting mechanisms.

While the later NH2-terminal cleavage events of proSP-C have been characterized in prior work (73) our focus here is on early trafficking through proximal compartments including the Golgi. Although this compartment is essential for proSP-C palmitoylation (74-78), it is also critical for initial proteolytic maturation and subsequent trafficking, highlighting the central role of classical anterograde transport patterns in complete SP-C biosynthetic maturation. In light of these observations, the fragmented morphology of the Golgi in I73T-expressing cells is particularly noteworthy. Although our current data do not resolve the temporal sequence between I73T SP-C mistrafficking and Golgi fragmentation, previous studies have shown that I73T expression induces metabolic reprogramming and a late-stage autophagy block (23, 27). The Golgi is increasingly recognized as a dynamic organelle that responds to such cellular stressors(79), including altered nutrient demands (80, 81) and autophagic flux (82-84). Notably, Golgi stacking proteins have been implicated in the regulation of autophagy and secretory pathways (80, 81, 85), further linking structural integrity to homeostatic functions.

In this context, our findings of fragmented Golgi architecture in I73T-expressing cells (**Figure 6**) suggest a model in which Golgi fragmentation may serve as a feedforward mechanism that exacerbates I73T mistrafficking. Specifically, fragmentation may impair the function of key resident Golgi enzymes involved in intra-Golgi substrate processing and accelerate cargo exit, as prior studies have shown that disruption of Golgi architecture can broadly impair sorting fidelity and compromise global trafficking efficiency (86, 87). Collectively, these data extend previous observations of post-Golgi dysfunction in I73T-expressing cells and highlight the interdependence of Golgi and endolysosomal compartments in maintaining epithelial proteostasis.

Importantly, arrest of protein transport in the Golgi caused accumulation of an intermediate proSP-C species, indicating that the earliest cleavage step occurs distal to the ER and prior to LRO (LB) delivery. Furthermore, this processing event was sensitive to DC1, a compound that inhibits PPCs (67) including Furin which was found to be highly enriched in AT2 cells (**Figure 7**). Bioinformatic analysis of the SP-C COOH-terminal propeptide revealed conserved multibasic recognition motifs used by PPCs, and site-directed mutagenesis of K160 and R167 attenuated proteolytic processing (**Figure S5)**. Notably, the location of this candidate convertase cleavage motif is compatible with our prior observation in primary AT2 cells using epitope specific antibodies that the first C-terminal cleavage of WT SP-C does not result in removal of the entire BRICHOS domain (5, 40, 60). Moreover, we note that this initial C-terminal cleavage is not unique to AT2 cells, as it is also observed in A549, HEK, and MLE-12 cells expressing SP-C^WT^. Our findings place SP-C within the broader BRICHOS protein family, several members of which, including Bri2 and GKN1, undergo PPC-dependent cleavage.

Several limitations should be considered when interpreting these results. Our in vitro cleavage assay uses a simplified, membrane-free system that reduces steric hindrance and lacks the membrane-associated context of the native secretory pathway, where factors such as subcellular localization and disulfide bonding may influence Furin activity. While our data implicate Furin in the initial C-terminal cleavage of SP-C, we do not exclude contributions from other Golgi-resident proteases or redundancy within the PPC family, as overlapping substrate motifs are common. Moreover, although many PPCs are enriched in the Golgi/TGN, several are known to recycle through the plasma membrane or endosomal compartments. Thus, we cannot rule out the possibility that a minor pool of SP-C if accessible as a substrate in these compartments may undergo cleavage, but our data suggest that the majority of this processing occurs within the late Golgi/TGN. Our study also does not address the precise mechanism by which the I73T mutant becomes mislocalized to the plasma membrane; however, we observe that preventing the initial C-terminal cleavage alters SP-C trafficking kinetics—delaying its exit from the Golgi and reducing steady-state colocalization with LAMP3⁺ compartments. Finally, we cannot exclude the presence of alternative or secondary cleavage sites within the BRICHOS domain that may be differentially recognized by other proteases. Despite these limitations, our findings offer new insights into the spatial and enzymatic regulation of early SP-C maturation and trafficking.

Taken together, our work defines a revised model of SP-C maturation in which the first COOH-terminal cleavage event is spatially and mechanistically linked to trafficking through the Golgi and is likely mediated by the PPC family member Furin. This study also helps to reconcile conflicting reports regarding proSP-C trafficking and emphasizes the need for physiologically relevant models—such as LRO-competent MLE-12 cells and iAT2 cells—when studying SP-C biosynthesis. In conclusion, our findings advance understanding of SP-C maturation by identifying a previously uncharacterized early proteolytic processing step shown to be essential to exclude aberrant transport of the proprotein to the plasma membrane, refining the model of SP-C trafficking, and highlighting the intricate regulation of protein processing in the Golgi – opening new avenues for investigating how misprocessing of SP-C might contribute to diseases such as idiopathic pulmonary fibrosis (IPF).

## AUTHOR CONTRIBUTIONS

MFB and SB developed the concept and designed experiments. SB, AR, and DJ performed experiments; LRR performed bioinformatic analysis. AR and SI generated cell lines and CC, RC, assisted with validation. CN performed immunogold EM; JK performed confocal microscopy on human lung explanted tissues. TEW provided samples from the SP-C^I73T^ ^KI/KI^ mouse. KDA and DNK generated iAT2s. YT performed animal studies. AM performed flow cytometry; SB and MFB analyzed data, generated figures, and interpreted results; MFB and SB drafted manuscript; MFB, SB, SM, KDA, DNK, and JAW edited the manuscript. All authors reviewed and approved the final version prior to submission.

## ACKNOWLEDGMENTS

Michael F. Beers is the Robert L. Mayock and David A. Cooper Professor of Medicine. We thank Jeremy Katzen for helpful discussions. We thank Didier Trono and David Root for plasmids and vectors.

## REFERENCES

1. Beers, M., and Mulugeta, S. (2005) Surfactant protein C biosynthesis and its emerging role in conformational lung disease Annual review of physiology 67,

2. Beers, M. F., and Fisher, A. B. (1992) Surfactant protein C: a review of its unique properties and metabolism. Am J Physiol (Lung Cell Mol Physiol) 263, L151–L160

3. Beers, M. F., and Moodley, Y. (2017) When Is an Alveolar Type 2 Cell an Alveolar Type 2 Cell? A Conundrum for Lung Stem Cell Biology and Regenerative Medicine Am J Respir Cell Mol Biol 57, 18–27

4. Beers, M. (1996) Inhibition of cellular processing of surfactant protein C by drugs affecting intracellular pH gradients The Journal of biological chemistry 271,

5. Johnson, A., Braidotti, P., Pietra, G., Russo, S., Kabore, A., Wang, W. et al. (2001) Post-translational processing of surfactant protein-C proprotein: targeting motifs in the NH(2)-terminal flanking domain are cleaved in late compartments American journal of respiratory cell and molecular biology 24,

6. Glasser, S. W., Korfhagen, T. R., Bruno, M. D., Dey, C., and Whitsett, J. A. (1990) Structure and expression of the pulmonary surfactant protein SP-C gene in the mouse. Journal of Biological Chemistry 265, 21986–21991

7. Beers, M. F., and Lomax, C. (1995) Synthesis and processing of hydrophobic surfactant protein C by isolated rat type II cells. Am J Physiol (Lung Cell Mol Physiol) 269, L744–L753

8. Mulugeta, S., and Beers, M. F. (2003) Processing of surfactant protein C requires a type II transmembrane topology directed by juxtamembrane positively charged residues Journal of Biological Chemistry 278, 47979–47986

9. Johnson, A. L., Braidotti, P., Pietra, G. G., Russo, S. J., Kabore, A., Wang, W. J. et al. (2001) Post-translational processing of surfactant protein-C proprotein - Targeting motifs in the NH2-terminal flanking domain are cleaved in late compartments American Journal of Respiratory Cell and Molecular Biology 24, 253–263

10. Kabore, A. F., Wang, W. J., Russo, S. J., and Beers, M. F. (2001) Biosynthesis of surfactant protein C: characterization of aggresome formation by EGFP chimeras containing propeptide mutants lacking conserved cysteine residues Journal of Cell Science 114, 293–302

11. Russo, S. J., Wang, W.-J., Lomax, C., and Beers, M. F. (1999) Structural requirements for intracellular targeting of surfactant protein C Am J Physiol (Lung Cell Mol Physiol) 277, L1034–L1044

12. Beers, M. F., Lomax, C. A., and Russo, S. J. (1998) Synthetic processing of surfactant protein C by alveolar epithelial cells. The COOH terminus of proSP-C is required for post-translational targeting and proteolysis Journal of Biological Chemistry 273, 15287–15293

13. Hamvas, A., Nogee, L., White, F., Schuler, P., Hackett, B., Huddleston, C. et al. (2004) Progressive lung disease and surfactant dysfunction with a deletion in surfactant protein C gene American journal of respiratory cell and molecular biology 30,

14. Nogee, L., Dunbar, A., Wert, S., Askin, F., Hamvas, A., and Whitsett, J. (2001) A mutation in the surfactant protein C gene associated with familial interstitial lung disease The New England journal of medicine 344,

15. Nogee, L., Dunbar, A., Wert, S., Askin, F., Hamvas, A., and Whitsett, J. (2002) Mutations in the surfactant protein C gene associated with interstitial lung disease Chest 121,

16. Abou, T. R., Jaubert, F., Emond, S., Le Bourgeois, M., Epaud, R., Karila, C., et al. (2009) Familial interstitial disease with I73T mutation: A mid- and long-term study Pediatric pulmonology 44,

17. Brasch, F., Griese, M., Tredano, M., Johnen, G., Ochs, M., Rieger, C., et al. (2004) Interstitial lung disease in a baby with a de novo mutation in the SFTPC gene The European respiratory journal 24,

18. Cameron, H., Somaschini, M., Carrera, P., Hamvas, A., Whitsett, J., Wert, S. et al. (2005) A common mutation in the surfactant protein C gene associated with lung disease The Journal of pediatrics 146,

19. van Moorsel, C., van Oosterhout, M., Barlo, N., de Jong, P., van der Vis, J., Ruven, H., et al. (2010) Surfactant protein C mutations are the basis of a significant portion of adult familial pulmonary fibrosis in a dutch cohort American journal of respiratory and critical care medicine 182,

20. Vorbroker, D. K., Voorhout, W. F., Weaver, T. E., and Whitsett, J. A. (1995) Posttranslational processing of surfactant protein C in rat type II cells Am J Physiol (Lung Cell Mol Physiol) 269, L727–L733

21. Voorhout, W. F., Veenendaal, T., Haagsman, H. P., Weaver, T. E., Whitsett, J. A., Van Golde, L. M., et al. (1992) Intracellular processing of pulmonary surfactant protein B in an endosomal/lysosomal compartment Am J Physiol (Lung Cell Mol Physiol) 263, L479-L486

22. Beers, M. F., Hawkins, A., Maguire, J. A., Kotorashvili, A., Zhao, M., Newitt, J. L. et al. (2011) A Nonaggregating Surfactant Protein C Mutant Is Misdirected to Early Endosomes and Disrupts Phospholipid Recycling Traffic 12, 1196-1210

23. Alysandratos, K.-D., Russo, S. J., Petcherski, A., Taddeo, E. P., Acín-Pérez, R., Villacorta-Martin, C., et al. (2021) Patient-specific iPSCs carrying an SFTPC mutation reveal the intrinsic alveolar epithelial dysfunction at the inception of interstitial lung disease Cell Reports 36, 109636

24. Maguire, J. A., Mulugeta, S., and Beers, M. F. (2011) Endoplasmic Reticulum Stress Induced by Surfactant Protein C BRICHOS Mutants Promotes Proinflammatory Signaling by Epithelial Cells American Journal of Respiratory Cell and Molecular Biology 44, 404–414

25. Maguire, J. A., Mulugeta, S., and Beers, M. F. (2012) Multiple ways to die: Delineation of the unfolded protein response and apoptosis induced by Surfactant Protein C BRICHOS mutants The International Journal of Biochemistry and Cell Biology 44, 101–112

26. Mulugeta, S., Maguire, J. A., Newitt, J. L., Russo, S. J., Kotorashvili, A., and Beers, M. F. (2007) Misfolded BRICHOS SP-C mutant proteins induce apoptosis via caspase-4- and cytochrome c-related mechanisms American Journal of Physiology-Lung Cellular and Molecular Physiology 293, L720–L729

27. Rodriguez, L. R., Alysandratos, K. D., Katzen, J., Murthy, A., Roque Barboza, W., Tomer, Y., et al. (2025) Impaired AMPK control of alveolar epithelial cell metabolism promotes pulmonary fibrosis JCI Insight 10,

28. Hawkins, A., Guttentag, S. H., Deterding, R., Funkhouser, W. K., Goralski, J. L., Chatterjee, S. et al. (2015) A non-BRICHOS SFTPC mutant (SP-CI73T) linked to interstitial lung disease promotes a late block in macroautophagy disrupting cellular proteostasis and mitophagy Am J Physiol Lung Cell Mol Physiol 308, L33–L47

29. Nureki, S. I., Tomer, Y., Venosa, A., Katzen, J., Russo, S. J., Jamil, S. et al. (2018) Expression of mutant Sftpc in murine alveolar epithelia drives spontaneous lung fibrosis The Journal of Clinical Investigation 128, 4008-4024

30. Katzen, J., Wagner, B. D., Venosa, A., Kopp, M., Tomer, Y., Russo, S. J., et al. (2019) An SFTPC BRICHOS mutant links epithelial ER stress and spontaneous lung fibrosis JCI Insight 4,

31. Rodriguez, L., Tomer, Y., Carson, P., Dimopoulos, T., Zhao, M., Chavez, K., et al. (2022) Chronic Expression of a Clinical SFTPC Mutation Causes Murine Lung Fibrosis with IPF Features Am J Respir Cell Mol Biol 10.1165/rcmb.2022-0203MA

32. Rodriguez, L. R., Tang, S. Y., Roque Barboza, W., Murthy, A., Tomer, Y., Cai, T. Q., et al. (2023) PGF2alpha signaling drives fibrotic remodeling and fibroblast population dynamics in mice JCI Insight 8,

33. Brasch, F., Ten Brinke, A., Johnen, G., Ochs, M., Kapp, N., Müller, K. et al. (2002) Involvement of cathepsin H in the processing of the hydrophobic surfactant-associated protein C in type II pneumocytes American journal of respiratory cell and molecular biology 26,

34. Kotorashvili, A., Mulugeta, S., Guttentag, S., and Beers, M. (2009) Pepsinogen C Is a Candidate Protease for Post-Translational Processing of Surfactant Protein C. In American Thoracic Society 2009 International Conference, May 15-20, 2009 • San Diego, California, American Thoracic Society, A6265

35. Dickens, J. A., Rutherford, E. N., Abreu, S., Chambers, J. E., Ellis, M. O., van Schadewijk, A., et al. (2021) Novel insights into surfactant protein C trafficking revealed through the study of a pathogenic mutant European Respiratory Journal 10.1183/13993003.00267-20212100267

36. Sitaraman, S., Na, C. L., Yang, L., Filuta, A., Bridges, J. P., and Weaver, T. E. (2019) Proteasome dysfunction in alveolar type 2 epithelial cells is associated with acute respiratory distress syndrome Sci Rep 9, 12509

37. Wikenheiser, K. A., Vorbroker, D. K., Rice, W. R., Clark, J. C., Bachurski, C. J., Oie, H. K. et al. (1993) Production of immortalized distal respiratory epithelial cell lines from surfactant protein C/simian virus 40 large tumor antigen transgenic mice Proceedings of the National Academy of Sciences of the United States of America 90, 11029–11033

38. Wang, W. J., Russo, S. J., Mulugeta, S., and Beers, M. F. (2002) Biosynthesis of Surfactant Protein C (SP-C). Sorting of SP-C proprotein involves homomeric association via a signal anchor domain Journal of Biological Chemistry 277, 19929–19937

39. Wang, W. J., Mulugeta, S., Russo, S. J., and Beers, M. F. (2003) Deletion of exon 4 from human surfactant protein C results in aggresome formation and generation of a dominant negative Journal of Cell Science 116, 683–692

40. Solarin, K. O., Ballard, P. L., Guttentag, S. H., Lomax, C. A., and Beers, M. F. (1997) Expression and glucocorticoid regulation of surfactant protein C in human fetal lung Pediatric Research 42, 356–364

41. Mulugeta, S., Nguyen, V., Russo, S. J., Muniswamy, M., and Beers, M. F. (2005) A Surfactant Protein C Precursor Protein BRICHOS Domain Mutation Causes Endoplasmic Reticulum Stress, Proteasome Dysfunction, and Caspase 3 Activation American Journal of Respiratory Cell and Molecular Biology 32, 521-530

42. Jacob, A., Morley, M., Hawkins, F., McCauley, K. B., Jean, J. C., Heins, H., et al. (2017) Differentiation of Human Pluripotent Stem Cells into Functional Lung Alveolar Epithelial Cells Cell Stem Cell 21, 472-488

43. Hawkins, F., Kramer, P., Jacob, A., Driver, I., Thomas, D. C., McCauley, K. B. et al. (2017) Prospective isolation of NKX2-1-expressing human lung progenitors derived from pluripotent stem cells J Clin Invest 127, 2277-2294

44. Jacob, A., Vedaie, M., Roberts, D. A., Thomas, D. C., Villacorta-Martin, C., Alysandratos, K.-D. et al. (2019) Derivation of self-renewing lung alveolar epithelial type II cells from human pluripotent stem cells Nature Protocols 14, 3303-3332

45. Tang, X., Snowball, J. M., Xu, Y., Na, C. L., Weaver, T. E., Clair, G. et al. (2017) EMC3 coordinates surfactant protein and lipid homeostasis required for respiration J Clin Invest 127, 4314-4325

46. Ridsdale, R., Na, C. L., Xu, Y., Greis, K. D., and Weaver, T. (2011) Comparative proteomic analysis of lung lamellar bodies and lysosome-related organelles PLoS One 6, e16482

47. Petrosyan, A., and Cheng, P. W. (2013) A non-enzymatic function of Golgi glycosyltransferases: mediation of Golgi fragmentation by interaction with non-muscle myosin IIA Glycobiology 23, 690–708

48. Venosa, A., Cowman, S., Katzen, J., Tomer, Y., Armstrong, B. S., Mulugeta, S., et al. (2021) Role of CCR2(+) Myeloid Cells in Inflammation Responses Driven by Expression of a Surfactant Protein-C Mutant in the Alveolar Epithelium Front Immunol 12, 665818

49. Venosa, A., Katzen, J., Tomer, Y., Kopp, M., Jamil, S., Russo, S. J. et al. (2019) Epithelial Expression of an Interstitial Lung Disease-Associated Mutation in Surfactant Protein-C Modulates Recruitment and Activation of Key Myeloid Cell Populations in Mice J Immunol 202, 2760-2771

50. Katzen, J., Rodriguez, L. R., Tomer, Y., Babu, A., Ming, Z., Murthy, A. et al. (2022) Disruption of Proteostasis Causes IRE1 Mediated Reprogramming of Alveolar Epithelial Cells Proceedings National Academy Sciences 119, e2123187119

51. Hoffman, E. T., Barboza, W. R., Rodriguez, L. R., Dherwani, R., Tomer, Y., Murthy, A., et al. (2024) Aberrant Transitional Alveolar Epithelial Cells Promote Pathogenic Activation of Lung Fibroblasts in Preclinical Fibrosis Modeling bioRxiv 10.1101/2024.06.17.5993512024.2006.2017.599351

52. Chen, S., Zhou, Y., Chen, Y., and Gu, J. (2018) fastp: an ultra-fast all-in-one FASTQ preprocessor Bioinformatics (Oxford, England) 34,

53. Langmead, B., and Salzberg, S. (2012) Fast gapped-read alignment with Bowtie 2 Nature methods 9,

54. Liao, Y., Smyth, G., and Shi, W. (2014) featureCounts: an efficient general purpose program for assigning sequence reads to genomic features Bioinformatics (Oxford, England) 30,

55. Law, C., Chen, Y., Shi, W., and Smyth, G. (2014) voom: Precision weights unlock linear model analysis tools for RNA-seq read counts Genome biology 15,

56. Liu, R., Holik, A., Su, S., Jansz, N., Chen, K., Leong, H., et al. (2015) Why weight? Modelling sample and observational level variability improves power in RNA-seq analyses Nucleic acids research 43,

57. Vorbroker, D. K., Dey, C., Weaver, T. E., and Whitsett, J. A. (1992) Surfactant protein C precursor is palmitoylated and associates with subcellular membranes Biochim Biophys Acta 1105, 161–169

58. Dutta, D., Williamson, C., Cole, N., and Donaldson, J. (2012) Pitstop 2 is a potent inhibitor of clathrin-independent endocytosis PloS one 7,

59. von Kleist, L., Stahlschmidt, W., Bulut, H., Gromova, K., Puchkov, D., Robertson, M., et al. (2011) Role of the clathrin terminal domain in regulating coated pit dynamics revealed by small molecule inhibition Cell 146,

60. Beers, M. F. (1996) Inhibition of cellular processing of surfactant protein C by drugs affecting intracellular pH gradients. Journal of Biological Chemistry 271, 14361–14370

61. Hirschberg, K., and Lippincott-Schwartz, J. (1999) Secretory pathway kinetics and in vivo analysis of protein traffic from the Golgi complex to the cell surface FASEB journal : official publication of the Federation of American Societies for Experimental Biology 13 Suppl 2,

62. Matlin, K., and Simons, K. (1983) Reduced temperature prevents transfer of a membrane glycoprotein to the cell surface but does not prevent terminal glycosylation Cell 34,

63. Saraste, J., and Kuismanen, E. (1984) Pre- and post-Golgi vacuoles operate in the transport of Semliki Forest virus membrane glycoproteins to the cell surface Cell 38,

64. Saraste, J., and Svensson, K. (1991) Distribution of the intermediate elements operating in ER to Golgi transport Journal of cell science 100 (Pt 3),

65. Willander, H., Hermansson, E., Johansson, J., and Presto, J. (2011) BRICHOS domain associated with lung fibrosis, dementia and cancer--a chaperone that prevents amyloid fibril formation? The FEBS journal 278,

66. Seidah, N., and Prat, A. (2012) The biology and therapeutic targeting of the proprotein convertases Nature reviews Drug discovery 11,

67. Komiyama, T., Coppola, J., Larsen, M., van Dort, M., Ross, B., Day, R. et al. (2009) Inhibition of furin/proprotein convertase-catalyzed surface and intracellular processing by small molecules The Journal of biological chemistry 284,

68. Nour, N., Basak, A., Chrétien, M., and Seidah, N. (2003) Structure-function analysis of the prosegment of the proprotein convertase PC5A The Journal of biological chemistry 278,

69. Duckert, P., Brunak, S., and Blom, N. (2004) Prediction of proprotein convertase cleavage sites Protein engineering, design & selection : PEDS 17,

70. Conkright, J. J., Bridges, J. P., Na, C. L., Voorhout, W. F., Trapnell, B., Glasser, S. W. et al. (2001) Secretion of surfactant protein C, an integral membrane protein, requires the N-terminal propeptide Journal of Biological Chemistry 276, 14658–14664

71. Kook, S., Wang, P., Meng, S., Jetter, C. S., Sucre, J. M. S., Benjamin, J. T., et al. (2021) AP-3- dependent targeting of flippase ATP8A1 to lamellar bodies suppresses activation of YAP in alveolar epithelial type 2 cells Proc Natl Acad Sci U S A 118,

72. Tobys, D., Kowalski, L., Cziudaj, E., Müller, S., Zentis, P., Pach, E., et al. (2021) Inhibition of clathrin-mediated endocytosis by knockdown of AP-2 leads to alterations in the plasma membrane proteome Traffic (Copenhagen, Denmark) 22,

73. Brasch, F., ten Brinke, A., Johnen, G., Ochs, M., Kapp, N., Muller, K. M. et al. (2002) Involvement of cathepsin H in the processing of the hydrophobic surfactant-associated protein C in type II pneumocytes American Journal of Respiratory Cell and Molecular Biology 26, 659–670

74. Flach, C. R., Gericke, A., Keough, K. M., and Mendelsohn, R. (1999) Palmitoylation of lung surfactant protein SP-C alters surface thermodynamics, but not protein secondary structure or orientation in 1,2-dipalmitoylphosphatidylcholine langmuir films Biochimica et Biophysica Acta 1416, 11–20

75. Kabore, A. F., Russo, S. J., Wang, W.-J., and Beers, M. F. (2000) Palmitoylation of proSP-C is not required for its intracellular targeting and proteolytic processing American Journal of Respiratory and Critical Care Medicine 161, A41

76. Na, N. P., Meyer, M. C., Flach, C. R., Mendelsohn, R., and Galla, H. J. (2007) Surfactant protein C and lung function: new insights into the role of alpha-helical length and palmitoylation Eur Biophys J 36, 477–489

77. Roldan, N., Goormaghtigh, E., Perez-Gil, J., and Garcia-Alvarez, B. (2015) Palmitoylation as a key factor to modulate SP-C-lipid interactions in lung surfactant membrane multilayers Biochim Biophys Acta 1848, 184–191

78. ten Brinke, A., van Golde, L. M. G., and Batenburg, J. J. (2002) Palmitoylation and processing of the lipopeptide surfactant protein C Biochimica et Biophysica Acta-Molecular and Cell Biology of Lipids 1583, 253–265

79. Geng, J., and Klionsky, D. J. (2010) The Golgi as a potential membrane source for autophagy Autophagy 6, 950–951

80. Zhang, X., Wang, L., Lak, B., Li, J., Jokitalo, E., and Wang, Y. (2018) GRASP55 Senses Glucose Deprivation through O-GlcNAcylation to Promote Autophagosome-Lysosome Fusion Dev Cell 45, 245–261 e246

81. Zhang, X., Wang, L., Ireland, S. C., Ahat, E., Li, J., Bekier, M. E., 2nd et al. (2019) GORASP2/GRASP55 collaborates with the PtdIns3K UVRAG complex to facilitate autophagosome-lysosome fusion Autophagy 15, 1787-1800

82. Takahashi, Y., Meyerkord, C. L., Hori, T., Runkle, K., Fox, T. E., Kester, M. et al. (2011) Bif-1 regulates Atg9 trafficking by mediating the fission of Golgi membranes during autophagy Autophagy 7, 61–73

83. Mattera, R., Park, S. Y., De Pace, R., Guardia, C. M., and Bonifacino, J. S. (2017) AP-4 mediates export of ATG9A from the trans-Golgi network to promote autophagosome formation Proc Natl Acad Sci U S A 114, E10697–E10706

84. Yamamoto, H., Kakuta, S., Watanabe, T. M., Kitamura, A., Sekito, T., Kondo-Kakuta, C. et al. (2012) Atg9 vesicles are an important membrane source during early steps of autophagosome formation J Cell Biol 198, 219–233

85. Nuchel, J., Tauber, M., Nolte, J. L., Morgelin, M., Turk, C., Eckes, B. et al. (2021) An mTORC1-GRASP55 signaling axis controls unconventional secretion to reshape the extracellular proteome upon stress Mol Cell 81, 3275-3293 e3212

86. Lee, I., Tiwari, N., Dunlop, M. H., Graham, M., Liu, X., and Rothman, J. E. (2014) Membrane adhesion dictates Golgi stacking and cisternal morphology Proc Natl Acad Sci U S A 111, 1849-1854

87. Puthenveedu, M. A., Bachert, C., Puri, S., Lanni, F., and Linstedt, A. D. (2006) GM130 and GRASP65- dependent lateral cisternal fusion allows uniform Golgi-enzyme distribution Nat Cell Biol 8, 238–248

