## Supplemental Data for "Divergent Pathways of Surfactant Protein C Maturation for Disease-Associated Isoforms"

**Running Title:** *Post-Translational Processing of ProSP-C Isoforms*

#### ¶ Correspondence should be addressed to:

**Michael F. Beers, M.D.**
Pulmonary and Critical Care Division
Perelman School of Medicine at The University of Pennsylvania
Edward J Stemmler Hall Suite 216
3450 Hamilton Walk
Philadelphia, Pennsylvania 19104-6118

### SUPPLEMENTAL FIGURES

#### FIGURES1

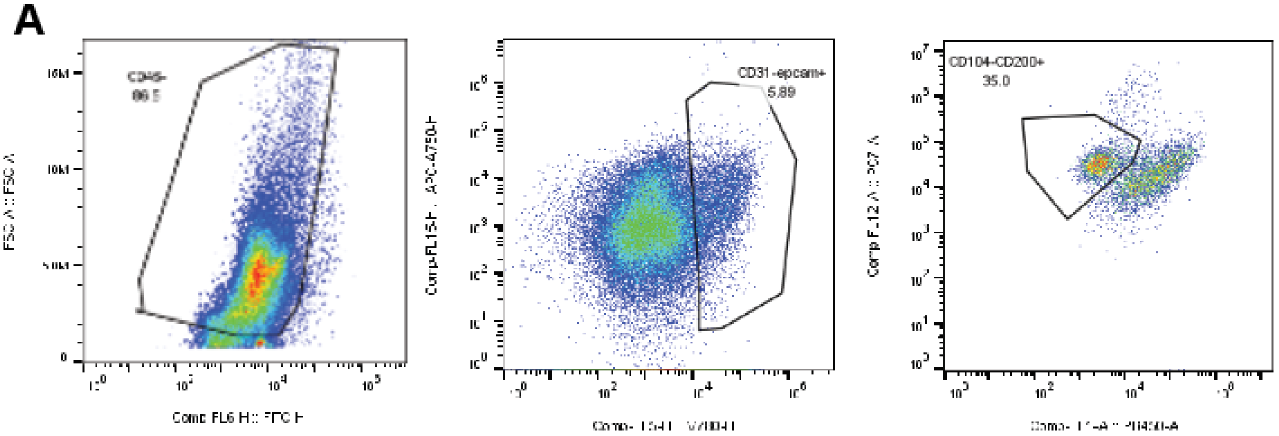

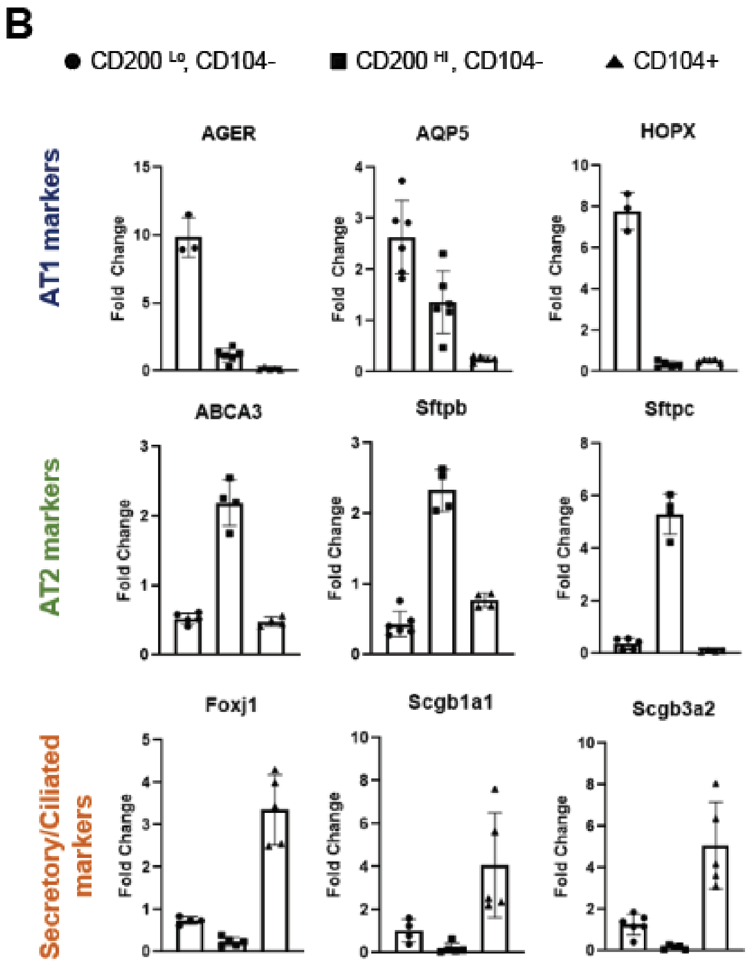

**Figure S1. Flow cytometry-based enrichment of AT2 cells and transcriptional validation of sorted epithelial populations. [A]** Representative gating strategy used to isolate lung epithelial cells and enrich for alveolar type 2 (AT2) cells. $CD45^{+}$ and $CD31^{+}$ cells were excluded, and epithelial subsets were further distinguished based on CD200 and CD104 surface expression. The $CD200^{Hi}$,$CD104^{-}$ population was found to be enriched for AT2 cells. **[B]** qRT-PCR analysis of sorted epithelial populations: $CD200^{Lo}$,$CD104^{-}$ (●), $CD200^{Hi}$,$CD104^{-}$ (■), and $CD104^{+}$ (▲). Expression of alveolar type 1 (AT1) markers (AGER, AQP5, HOPX), AT2 markers (ABCA3, SFTPB, SFTPC, and secretory/ciliated cell markers (FOXJ1, SCGB1A1, SCGB3A2) validates the enrichment of sorted populations. Data are shown as fold change relative to the $CD200^{Lo}$,$CD104^{-}$ group. Bars represent mean ± SEM.

FIGURES2

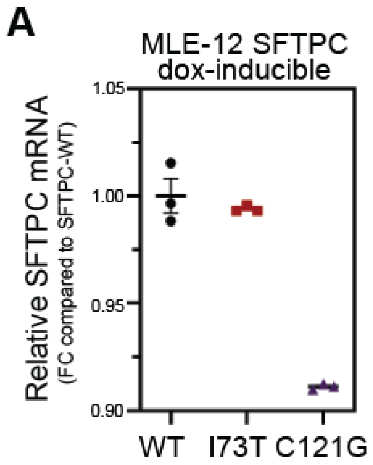

**Figure S2. SFTPC transcript levels in dox-inducible MLE-12 cells.**
**[A]** Quantitative RT-PCR analysis of SFTPC mRNA expression in MLE-12 cells stably expressing wild-type (WT), I73T, or C121G SFTPC following 24 hours of doxycycline induction (2.5 μM). Transcript levels are shown relative to WT and normalized to 18S.

#### FIGURES3

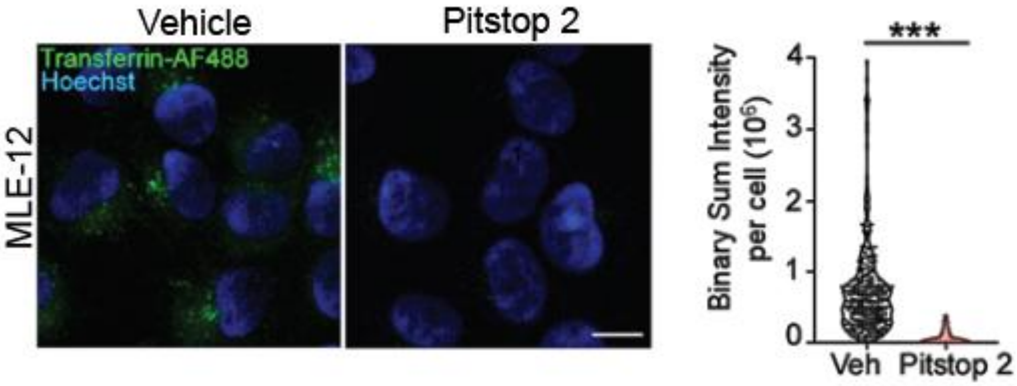

**Figure S3. Pitstop 2 treatment inhibits clathrin-mediated endocytosis in MLE-12 cells.**
**[A]** MLE-12 cells were preincubated with either DMSO (vehicle control) or 20 μM Pitstop 2 for 20 minutes at 37°C. Following preincubation, Alexa Fluor 488-conjugated transferrin (green) was added for 30 minutes at 37°C to allow for clathrin-mediated endocytosis in the continued presence of DMSO or Pitstop 2. Cells were then washed to remove surface-bound transferrin, fixed, and imaged to measure the pool of internalized transferrin. Nuclei were counterstained with Hoechst (blue). Representative images show robust transferrin uptake in vehicle-treated cells and markedly reduced uptake in Pitstop 2-treated cells. **[B]** Quantification (right) of binary sum intensity per cell confirms significant reduction in transferrin uptake following Pitstop 2 treatment (***$p < 0.001$). Scale bar: 10 μm.

**FIGURES4**

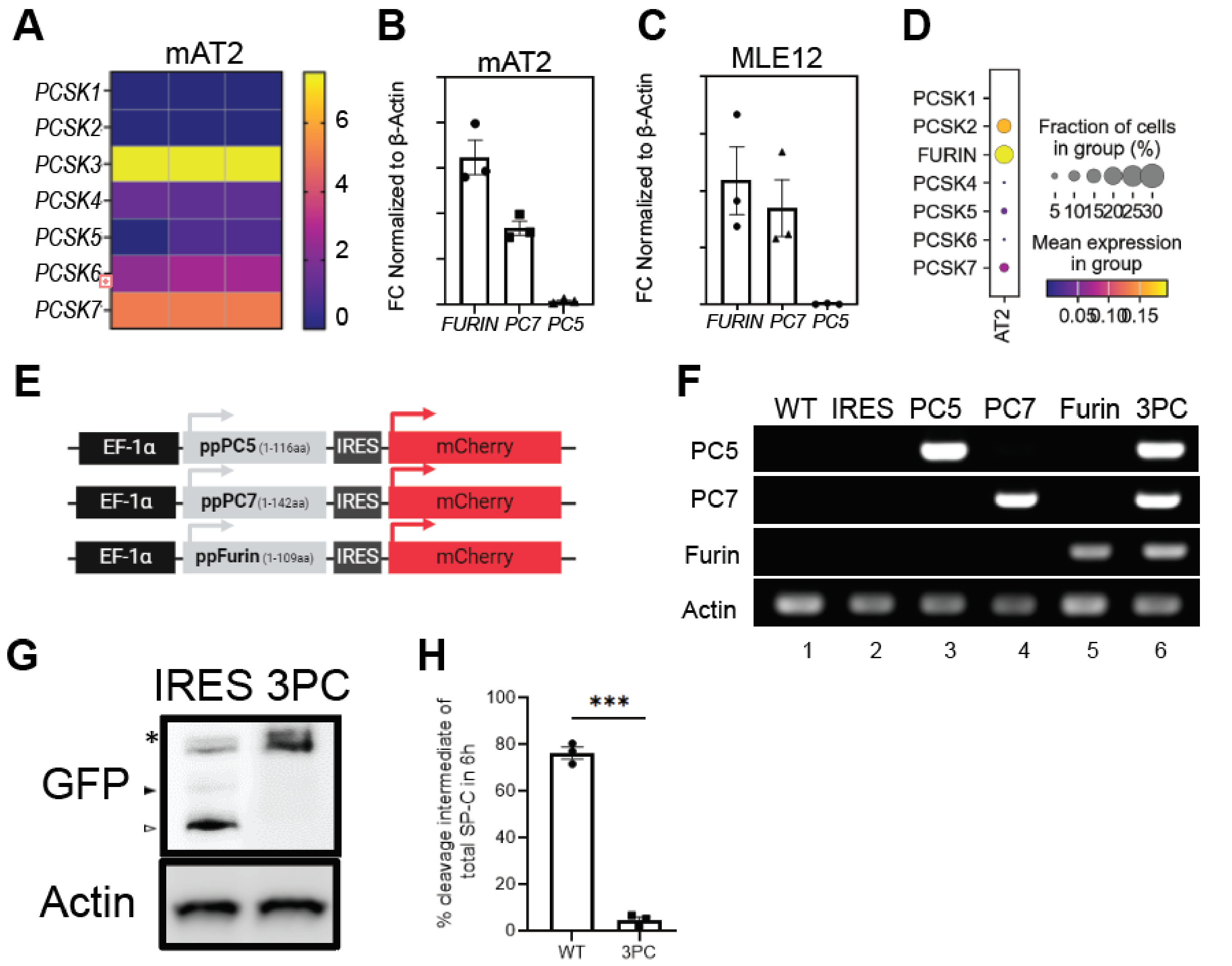

**Figure S4. Evaluation of candidate proprotein convertases involved in SP-C cleavage**. **[A]** Heatmap showing expression proprotein convertase family members (Pcsk1–Pcsk7) in primary mouse alveolar type 2 (mAT2) cells. **[B-C]** qRT-PCR analysis of *Pcsk3 (Furin), Pcsk7, and Pcsk5* transcript levels in mAT2 cells and MLE-12 cells normalized to β-actin. **[D]** Dot plot showing expression of *PCSK* genes in human AT2 cells. **[E]** Schematic of lentiviral constructs used to express N-terminal pre-prosegment of proprotein convertases (PC5, PC7, Furin) under EF-1α promoter, followed by IRES and mCherry. These truncated forms have been shown to inhibit**. [F]** RT-PCR validation of PC5, PC7, and Furin expression in MLE-12 cells transduced with lentiviral vectors. WT and IRES-only controls shown in lanes 1 and 2, respectively. Successful expression of pre-prosegments of individual proprotein convertases (lanes 3–5) and all three in combination (3PC, lane 6) confirmed in established cell lines. **[G]** Western blot of MLE12 cell line expressing all three inhibitory pre-proprotein convertase fragments (3PC) were induced with doxycycline for 6h and whole cell lysates were assessed via western blot to determine SP-C c—term cleavage (arrowheads). **[H]** Quantification of percent of SP-C cleaved intermediate in each condition.

#### FIGURES5

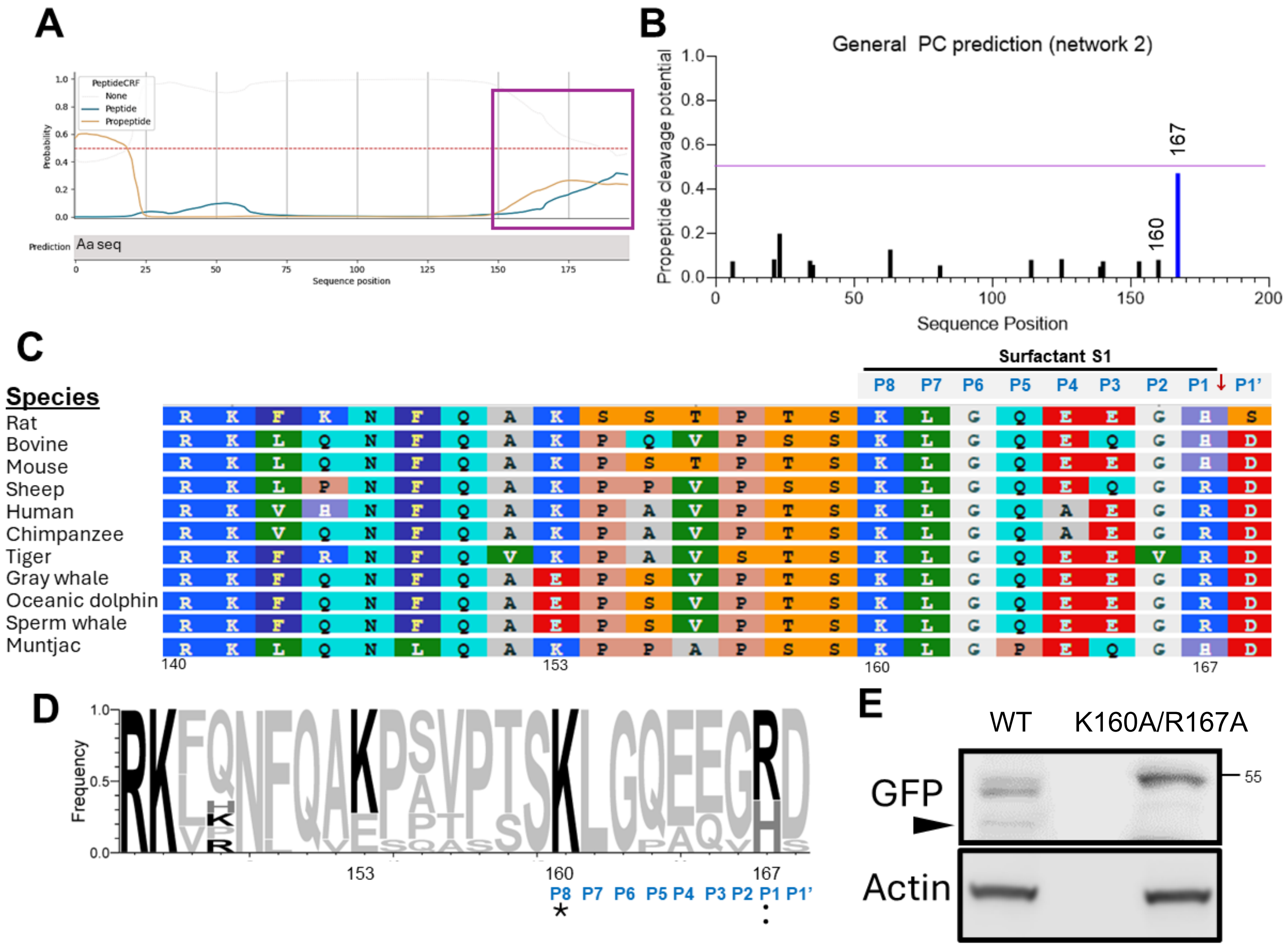

**Figure S5. Cleavage of WT proSP-C occurs within the BRICHOS domain of the COOH-propeptide.**
**[A]** DeepPeptide analysis of hSP-C predicts a C-terminal cleaved peptide around aa 150-197. https://biolib.com/KU/DeepPeptide **[B]** ProP 1.0 server predicts cleavage sites in hSP-C with the highest scoring site at aa167. **[C]** Sequence alignment across various species shows high degree of conservation. Sequence identification row (blue font): basic residue positions in the S1 cleavage site from P8 to P1′. Red arrow indicates the predicted site of furin cleavage. **[D]** To visualize the diversity of residues at each position of the S1 site, sequences were subjected to WebLogo 3.1 analysis (http://weblogo.threeplusone.com/create.cgi) with the frequency of residue found at each position displayed. **[E]** MLE12 transfected with GFP-SPC mutated at K160A/R167A shows diminished C-term cleavage processing (arrowhead) as compared to GFP-$SPC^{WT}$.

### TABLE 1 ANTIBODIES

| Antibodies | SOURCE | IDENTIFIER |
|---|---|---|
| Beta Actin | Proteintech | HRP-60008, AB_28/19183 |
| GAPDH | Proteintech | HRP-60004, RRID:AB_2737588 |
| GFP | Proteintech | |
| E-Cadherin | BD Biosciences | 610181 |
| GM130 | BD Biosciences | 610822 |
| TGN46 | ProteinTech | 13573-1-AP |
| Calreticulin | ProteinTech | 27298-1-AP |
| LAMP1 | DSHB | H4A3 |
| TOM20 | Cell Signaling | 42406 |
| Furin | Santa Cruz | sc-133142 |
| Furin | ProteinTech | 18413-1-AP |
| LAMP3 | Sant Cruz | Sc-5275 |
| HK1 | Proteintech | 19662-1-AP |
| SQSTM1/p62 | Cell Signaling | 5114 |
| LC3B | Cell Signaling | 2775S; RRID:AB_915950 |
| Pro SP-C (N terminal) | Beers et al. 1994 | N/A |
| Pro SP-B (PT3) | Beers et al. 1992 | N/A |
| Goat Anti-Mouse IgG (H+L), HRP conjugated | Biorad | 0300-0108P; RRID:AB_808614 |
| Goat Anti-Rabbit IgG (H+L), HRP conjugated | Biorad | 5196-2504; RRID:AB_619908 |
| ABCA3 Mouse | Seven Hills Bioreagents | RRID:AB_577285 |
| ABCA3 Rabbit | Seven Hills Bioreagents | RRID: AB_3146162 |
| SFTPC Rabbit N-term | Seven Hills Bioreagents | RRID:AB_2335890 |
| SFTPC Guinea pig | In house | |
| ***Flow cytometry Antibodies*** | | |
| CD45 (30F-11) BB515 | Biolegend | 564590 |
| Epcam (G8.8) BV711 | Biolegend | 118233 |
| Epcam (G8.8) BV785 | BD Biosciences | 118245 |
| CD31 (MEC13.3) APC/Cy7 | Biolegend | 102533 |
| CD104 (346-11A) PE/Cy7 | Biolegend | 123615 |
| CD51 (RMV-7) PE | Biolegend | 104105 |
| ***Cell stains*** | | |
| Lysotracker™ Deep Red | Invitrogen | L12492 |
| WGA AlexaFluor594 | Invitrogen | W11262 |
| ER-Tracker™ Red | Thermofisher | E34250 |

### TABLE 2 PRIMERS

| Oligonucleotides | | |
|---|---|---|
| Abca3 | Thermofisher | Mm01299912_m1 |
| Sftpb | Thermofisher | Mm00455678_m1 |
| Sftpc | Thermofisher | Mm00488144_m1 |
| Mouse_Ager_Forward (CTTGCTCTATGGGGAGCTGTA) | Sigma | |
| Mouse_Ager_ Reverse (GGAGGATTTGAGCCACGCT) | Sigma | |
| Mouse_Aqp5_Forward (TCTTGTGGGGATCTACTTCACC) | Sigma | |
| Mouse_Aqp5_ Reverse (TGAGAGGGGCTGAACCGAT) | Sigma | |
| Mouse_Scgb3a2_Forward (AGAAGTGTGTGGACGAGCTG) | Sigma | |
| Mouse_Scgb3a2_ Reverse (CAGGTGTGAAAGAGCCTCAAATG) | Sigma | |
| Mouse_Scgb1a1_Forward (AACATCATGAAGCTCACGGAGA) | Sigma | |
| Mouse_Scgb1a1_ Reverse (AGGGCAGTGACAAGGCTTTA) | Sigma | |
| Mouse_Foxj1_Forward (GGGAGGTGGGAGGAACTTCT) | Sigma | |
| Mouse_Foxj1_ Reverse (CGAATGTGAGGCCTGGCT) | Sigma | |
| ***Cloning oligos*** | | |
| NotI-ppFurin-For (CTAGCGGCCGCgccaccATGGAGCTGAGATCCTGGTTGC) | IDT | |
| BamHI-ppFurin-Rev (TCAGGATCCTTACACGTCCCTCTTGGCTCTTCG) | IDT | |
| ppPC7 (CTAGCGGCCGCgccaccATGCCGAAAGGGAGGCAGAAAGTCCCACACTTGGATGCCCACCTGGGCCTGCCCATCTGCCTCTGGCTGGAATTAGCCATCTTCTTTCTGGTTCCCCAGGTCATGGGCCTATCAGAGGCAGGTGGGCTTGACATCTTGGGCACAGGGGGGCTGAGCTGGGCCGTACATCTGGACAGCCTAGAAGGTGAGAGGAAGGAAGAGAGTCTG) | IDT gBlock | |
| ppPC5 (CTAGCGGCCGCgccaccATGGACTGGGACTGGGGGAACCGCTGCAGCCGCCCGGGACGGCGGGACCTGCTGTGCGTGCTGGCACTGCTCGCCGGCTGTCTGCTCCCGGTATGCCGGACGCGCGTCTACACCAACCACTGGGCAGTGAAGATCGCCGGCGGCTTCGCGGAGGCAGATCGCATAGCCAGCAAGTACGGATTCATCAACGTAGGACAGATCGGTGCA) | IDT gBlock | |
